# A PKM2-YAP reciprocal repartitioning modulates invasion of breast cancer cells

**DOI:** 10.1101/2025.07.10.664113

**Authors:** Mallar Banerjee, Mayank Pandhari, Sudiksha Mishra, Kottpalli Vidhipriya, Ramray Bhat

**Author notes:** Correspondence: Ramray Bhat.

## Abstract

Invasive cancer cells exhibit distinct morphomigrational and metabolic traits when confronted with biophysically variant matrix microenvironments enroute metastasis. Whether dynamical shifts in such traits are interlinked through a common molecular program remains ill-understood. Using triple negative breast cancer cell lines on Collagen I substrata coated hydrogels recapitulating stiffness values of non-cancerous breast tissue and the desmoplasia of tumors, we observed greater cell shape polarization and migration in the latter. Associated lower lactate and pyruvate levels in such conditions motivated us to examine their pyruvate kinase M2 expression, which showed nuclear and cytoplasmic localization in softer and stiffer environments respectively. Pharmacologically impairing PKM2 activity in stiffer substrata decreased migration and shape polarization of cancer cells while increasing lactate and pyruvate levels. In contrast, increasing its activity on softer substrata attenuated cancer cell migration and elongation. We assayed for localization of the mechanosensory protein YAP upon PKM2 activity modulation: PKM2 activation increased nuclear YAP localization on soft substrata. Pharmacologically inhibiting YAP on stiff substrata not just decreased migration but also increased nuclear localization of PKM2 and lactate and pyruvate levels. We propose that a reciprocal repartitioning of PKM2-YAP interlinks the cognate metabolic and migrational states of cancer cells; targeting such positive feedback may hold the key to future therapeutic strategies.

## Introduction

Invasion of cancer is an emergent outcome of dynamic and reciprocal cues being transduced by transformed cells from their surrounding extracellular matrix (ECM) (Pally, Goutham, et al., 2022; Sleeboom et al., 2024; Winkler et al., 2020). Such cues maybe biochemical, architectural or biophysical in nature (So & Tanner, 2021; Tien et al., 2020). The viscoelastic nature of the polymeric ECM entails that cancer cells are subject to, and respond to, the stiffness of the ECM that they confront as they migrate from their primary focus (Mierke, 2021). Stiffness of the matrix microenvironment is not only among the oldest changes that are relevant to diagnosis, it also an exemplar of how cancer remodels its surroundings during migration (Baker et al., 2009).

A plethora of studies in the last two decades have elegantly demonstrated how the mechanical properties of the ECM surrounding cancer cells play crucial roles in regulating proliferation, motility, shape polarization, and refractoriness to chemotherapy (Acerbi et al., 2015; Ishihara & Haga, 2022; Li et al., 2023; Xu et al., 2023) Experiments across different cancer types demonstrate that epithelial cells cultured on soft (low elastic modulus) ECM retain an epithelioid morphology, whereas those cultured on stiff (high elastic modulus) ECM exhibit front-back polarization and migrate faster (Wozniak et al., 2003). Transcriptomic and signalling experiments have been conducted in such variant microenvironments and the Hippo pathway and its constituent transcriptional regulator YAP has been identified as a canonical transducer of extracellular rheological cues (Ishihara & Haga, 2022). YAP is localized to the nucleus when cells are cultured on stiff substrata leading to transcription of pro-proliferative and anti-apoptotic genes, thus regulating cancer migration (W. Chen et al., 2021; Ishihara & Haga, 2022; Wang et al., 2024). This is an example of a mechanosensing - a trait by which cancer cells sense, re-adjust and respond to the biophysical variation in their surroundings.

Investigations in cancer invasion have also focused on elucidating the changes in metabolic dynamics upon cell transformation. While Warburg’s seminal observations established the increased reliance of cancer cells on aerobic glycolysis (Warburg et al., 1927), recent research shows that cancer cells can dynamically switch between glycolysis and mitochondrial oxidative phosphorylation or even sustain both at high levels depending on the context (DeBerardinis & Chandel, 2020; Evans et al., 2021; Hensley et al., 2016; Uslu et al., 2025). In high density Collagen I, Morris and coworkers report a lower reliance of murine 4T1 mammary cancer cells on glucose and a higher reliance on glutamine for oxidative phosphorylation (Morris et al., 2016). This metabolic plasticity is essential to progression as it feeds the anabolic demands for mitotic synthesis as well as the energetic demands for motility. Further exemplification of the plasticity is demonstrated through observations that highly migrating ‘leader’ cells in invading breast cancer masses selectively exhibit an increased glucose uptake (J. Zhang et al., 2019). Conversely, Commander and coworkers reported that in certain lung and breast cancer cell lines, cells at the invasive edge preferentially utilize oxidative phosphorylation, with ‘follower’ cells instead reliant on glycolysis (Commander et al., 2020). These findings underscore the heterogeneity of metabolic strategies employed by cancer cells during migration and invasion.

Can metabolic flux act as a regulator of the kinetics in cell phenotypic trait readjustments when confronted by variant rheological cues in the microenvironment? Modern efforts have begun to investigate this question with one particular study demonstrating how mechanical cues drive the actin to stabilize a rate limiting enzyme of glycolysis phosphofructokinase, resulting in higher glucose uptake and ATP production (Park et al., 2020). Using quantitative mass spectrometry, Goransson and coworkers have also recently shown how stiffness reprograms sterol/isoprenoid metabolism in breast cancer cells in a Rac1-dependent manner (Göransson et al., 2025). Even so, mechanistic links between the canonical mechanosensing pathways and metabolic dynamics in migrating cancer cells remain elusive (Parlani et al., 2022).

To address this lacuna, we were motivated by a tantalizing observation specific to another enzyme of glycolysis: pyruvate kinase M2 (PKM2), which catalyzes the final rate limiting step of the conversion of phosphoenolpyruvate to pyruvate. PKM2 is known to shuttle between cytoplasm and nucleus suggesting a temporally rapid mechanism by which a cell can utilize the same protein for distinct functions (Z. Zhang et al., 2019). Indeed, PKM2 exists in tetrameric and dimeric forms: the tetrameric form is catalytically active in the cytoplasm driving pyruvate production at optimal levels, whereas the dimeric form localizes to the nucleus, where it acts as a transcriptional co-activator of c-Myc to induce aerobic glycolysis and promote tumor progression (Amin et al., 2019; Christofk et al., 2008; Gao et al., 2012; Yang et al., 2011). Recent work by Liang and colleagues demonstrated that nuclear delocalization and tetramer formation of PKM2 are critical for non-small cell lung cancer progression (L. J. Liang et al., 2024).

In the present study, we use Collagen I-coated polyacrylamide gels with elastic moduli recapitulating the extremes of what is seen in the breast tissue (0.4 kPa in untransformed regions and 20 kPa, representing the desmoplastic zones) (Ishihara & Haga, 2022). Cultivating aggressive triple negative breast cancer cells on such matrices, we observed cytoplasmic and nuclear localization of PKM2 in stiffer and softer substrate respectively with greater shape polarisation and migration in stiffer substrata. Pharmacological modulation of PKM2 not just reverses the associated morphomigrational features of cancer cells but also the subcellular localization of YAP. The reciprocity of such effects with YAP modulation and the effect of the latter on lactate levels reveals the deep integration of mechanosensing and metabolic reprogramming that drives cancer invasion and suggests novel targets for management in the future.

## Results

### Matrix stiffness regulates shape polarization and migration dynamics of breast cancer cells

Previous studies have shown that increasing the stiffness of ECM alters the morphology and increases migrative capacity of cancer cells (Acerbi et al., 2015; Xu et al., 2023)). We began our studies by validating these observations using Collagen I-coated PA hydrogels of elastic moduli 0.4 kPa which represents mechanically the softest substrata within the transformed breast microenvironment (typifying non-cancerous breast tissues), and 20 kPa, which represents the stiffness of desmoplastic regions within the tumors. We assayed the morphological alterations of triple-negative breast cancer (TNBC) cell lines MDA-MB-231 and HCC1806 cultured on these scaffolds. Both cancer cells showed a relatively circular shape on 0.4 kPa hydrogels, whereas on 20 kPa hydrogels, the cells were more elongated (Figure 1A for MDA-MB-231 and S1B for HCC-1608; white signal represents DNA stained by DAPI and green represents F-actin stained by phalloidin). The mean cell area was found to be higher on the stiffer Collagen I coated substrata, and their aspect ratio was significantly reduced, when compared with the softer gels confirming higher spreading and greater spindle shape of cells on stiffer substrata (Figure 1A right depicting graphs for cell area and aspect ratio for MDA-MB-231 and Figure S1B right showing changes in cell area and aspect ratio for HCC-1608, p < 0.0001 for cell area and aspect ratio; significance assessed using unpaired Students t-test). We next assessed the motility of cancer cells seeded on soft and stiff scaffolds through timelapse videography for 3 h at an interval of 3 min/image acquisition. Consistent with previous studies, our results also indicated that migration speeds were elevated on stiff substrata compared with their softer counterparts (Figure 1B for MDA-MB-231 and S1C for HCC-1608; migration tracked for RFP-expressing MDA-MB-231 and GFP-expressing HCC1806 cells with individual tracks shown in distinct colors; graph showing significantly higher mean migration speeds on 20 kPa; p < 0.0001; significance assessed using unpaired Students t-test; video S1 for MDA-MB-231 on 0.4 kPa hydrogel, video S2 for MDA-MB-231 cells on 20 kPa hydrogels, video S9 for HCC1806 cells on 0.4 kPa hydrogels and video S10 for HCC1806 cells on 20 kPa hydrogels respectively). Cell shape change and motility modulation require spatially segregated F-actin polymerization-depolymerization, which expends ATP through a change in metabolic dynamics (Bays et al., 2017; Doostmohammadi & Ladoux, 2022; Olson et al., 2008; Park et al., 2020). We therefore assayed for levels of lactate and pyruvate to assess the glucose catabolic flux within cells as a function of stiffness. On softer gels, MDA-MB-231 cells showed higher extracellular and intracellular levels of lactate and pyruvate compared with cells on stiffer gels (Figure 1C; p < 0.0001, p < 0.0001, p = 0.008 and p = 0.001 for extracellular and intracellular lactate and pyruvate respectively, significance assessed using unpaired Students t-test). This suggested that in softer milieu, cancer cells convert pyruvate to lactate, both of which are secreted to the extracellular surroundings, whereas in stiffer environments, highly deformable and motile cancer cells may preferentially funnel glycolytic intermediates into other pathways, in the process depleting them. This led us to examine the role of pyruvate kinase M2, (PKM2), which is elevated in breast cancer progression (Figure S1D) (Chandrashekar et al., 2022; X. Chen et al., 2020)and whose localization has been recently demonstrated to play critical roles in modulating glycolytic flux (Amin et al., 2019). Through fluorescent immunocytochemistry, we observed that PKM2 was predominantly localized in the nucleus of cancer cells on softer substrata, whereas on stiffer substrata, it was localized predominantly in the cytoplasm (Figure 1D for MDA-MB-231 and S1E for HCC-1608; white signal represents DNA stained by DAPI and red represents cognate antibody signals; stiffness-cognate nucleocytoplasmic ratio of PKM2 shown in graph on the right, p < 0.0001; significance assessed using unpaired Student’s t-test). The association between stiffness-dependent trait alterations and PKM2 subcellular localization is depicted in Figure 1E. We next sought to test whether the activity of PKM2 regulated cellular morphomigrational behavior that was contextual to ECM stiffness.

**Figure 1.**
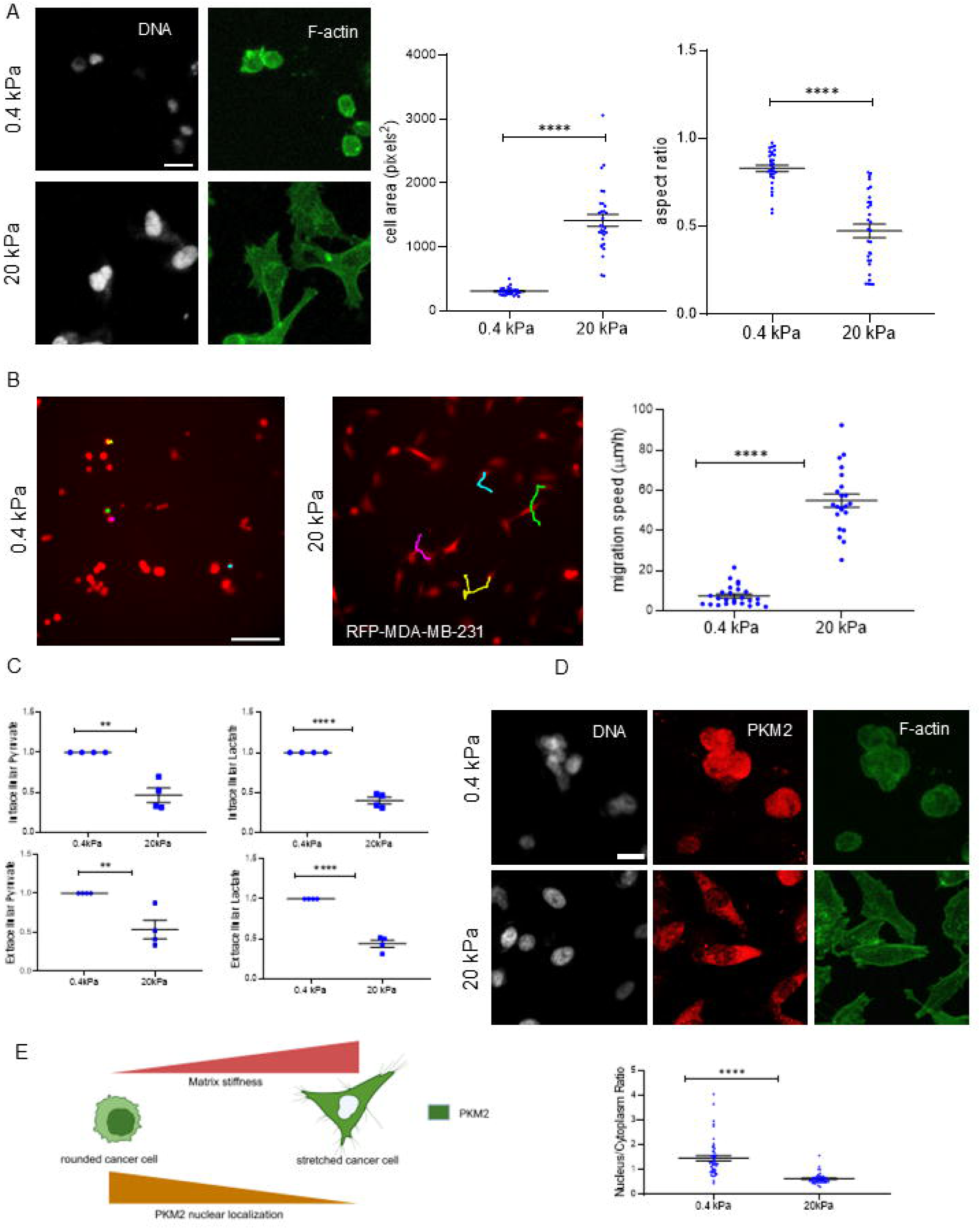
Matrix stiffness regulates morpho-migration dynamics of and subcellular PKM2 localization of MDA-MB-231 breast cancer cells. (A) Representative confocal photomicrographs of breast cancer MDA-MB-231 cells grown on 0.4 kPa (top row of micrographs) and 20 kPa (bottom row of micrographs) polyacrylamide hydrogels coated with Collagen I and observed after 24Lh of culture with staining for DNA (white using DAPI, left panel) and F-actin (green using phalloidin, right panel (n□= 3). Scale bar: 20□μm and graphs showing cell area (left) and aspect ratio (right) of individual MDA-MB-231 cells grown on 0.4 kPa and 20 kPa hydrogels (n□= 3, N□= 30 single cells). (B) Epifluorescence photomicrographs of RFP-labelled MDA-MB-231 cells with their migration tracks for 3 hours, 0.4 kPa (left) and 20 kPa (right) and scatter plot graph depicting migration speed (n=3, N=20). See also Video S1, S2. (C) Graph measuring fold change of extracellular lactate (top left), extracellular pyruvate (top right), intracellular lactate (bottom left), intracellular pyruvate (bottom right) between 0.4 kPa and 20 kPa hydrogels. (D) Representative confocal photomicrographs of MDA-MB-231 cells grown on 0.4 kPa (top row of micrographs) and 20 kPa (bottom row of micrographs) polyacrylamide hydrogels coated with Collagen I and observed after 24□h of culture with staining for DNA (white using DAPI, left panel), PKM2 (Red, middle panel) and F-actin (green, right panel). (n□= 3). Scale bar: 20□μm and scatter plot graph depicting nucleus/cytoplasm ratio of PKM2 (n=3, N=30) (See Figure S1 for a schematic depiction of how nuclear and cytoplasmic boundaries were delineated and the ratio of the PKM2 levels calculated). (E) Schematic depiction of matrix stiffness-driven phenotypic changes and PKM2 subcellular localization. Error bars denote mean□±□SEM. Unpaired Student’s□*t*□test was performed for statistical significance (\**P*□≤ 0.05, \*\**P*□≤ 0.01, \*\*\**P*□< 0.001, \*\*\*\**P*□< 0.0001).

### PKM2 cytoplasmic activation increases cancer migration on soft matrices

We treated MDA-MB-231 and HCC1806 cells with TEPP-46, a PKM2 cytoplasmic activator to increase cytoplasmic localization of PKM2 in cancer cells cultured on soft substrata (Anastasiou et al., 2012; Jiang et al., 2010). We observed cytoplasmic localization of PKM2 of cells on soft gels using 40 µm of TEPP-46 (Figure 2A for MDA-MB-231 and, S2A for HCC-1608; white signal represents DNA stained by DAPI and red represents cognate antibody signals). This was also confirmed through an assessment of the nuclear/cytoplasmic ratio of PKM2 which was found to be reduced after TEPP-46 treatment (Figure 2A right for MDA-MB-231, p < 0.0001 and Figure S2A right for HCC1806 with p = 0.0099; significance assessed using unpaired Student’s t-test). Cytoplasmic localization of PKM2 in MDA-MB-231 was found to increase mean cell area and to decrease aspect ratio indicating higher elongation and cell spreading (Figure 2B with white signal represents DNA stained by DAPI and green represents F-actin stained by phalloidin, p < 0.001 for both cell area and aspect ratio) on soft matrices. HCC1806 cells showed higher mean cell area indicating greater spreading upon cytoplasmic localisation of PKM2, whereas aspect ratio was not altered (Figure S2B with white signal represents DNA stained by DAPI and green represents F-actin stained by phalloidin, p < 0.001 for total cell area. Insignificant alteration in aspect ratio of HCC1806 cells could possibly be explained by the fact that the cell line is epithelioid in nature). Furthermore, to decipher if PKM2 activation modulates cell migration, timelapse imaging was performed. PKM2 activation significantly increased cancer cell migration on soft gels relative to controls (Figure 2C for MDA-MB-231 and S2C for HCC-1608; migration tracked for RFP-expressing MDA-MB-231 and GFP-expressing HCC1806 cells with individual tracks shown in distinct colors; video S3, video S4 for MDA-MB-231 and video S11, video S12 for HCC1806). These results suggest that increasing the cytoplasmic activity of PKM2 on softer substrata phenocopied their morphomigrationary behavior on stiffer ECM (see schematic depiction of PKM2 activation-driven phenotypic changes in Figure 2D).

**Figure 2.**
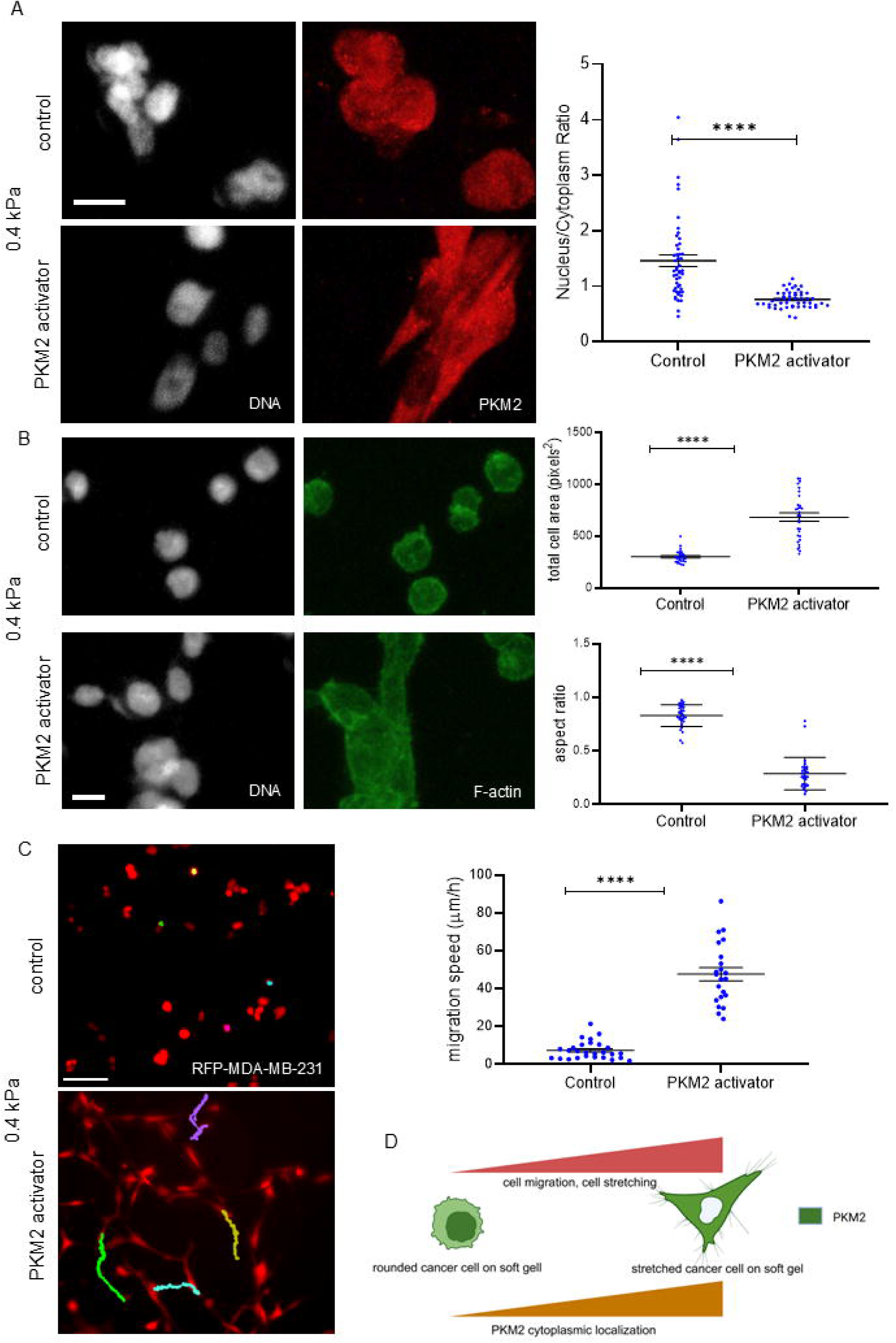
MDA-MB-231 breast cancer cell migration increases on soft matrices upon cytoplasmic PKM2 activation. (A) Representative confocal photomicrographs of breast cancer MDA-MB-231 cells untreated (top row of micrographs) or treated with 40□μM TEPP-46, PKM2 cytoplasmic activator (second row from top) grown on 0.4 kPa (soft) hydrogels coated with Collagen I and observed after 24□h of culture with staining by□DNA (white using DAPI, left panel) and PKM2 (Red, right panel). (n□= 3). Scale bar: 20□μm. Scatter plot graph (right) depicting nucleus/cytoplasm ratio of PKM2 between untreated and 40□μM TEPP-46-treated MDA-MB-231 cells on 0.4 kPa hydrogels (n = 3, N = 30). (B) Representative confocal photomicrographs of breast cancer MDA-MB-231 cells untreated (top row of micrographs) or treated with 40□μM TEPP-46, PKM2 cytoplasmic activator (second row from top) grown on 0.4 kPa (soft) hydrogels coated with Collagen I and observed after 24□h of culture with staining by□DNA (white using DAPI, left panel) and F- actin (green using phalloidin, right panel (n□= 3). Scale bar: 20□μm. Graphs showing cell area (top) and aspect ratio (bottom) of individual MDA-MB-231 cells untreated and treated with 40□μM TEPP-46 on 0.4 kPa hydrogels (n□= 3, N□= 30 single cells). (C) Epifluorescence photomicrographs of RFP-labelled MDA-MB-231 cells with their migration tracks for 3 hours, control (top) and treated with 40□μM TEPP-46 (bottom) on 0.4 kPa hydrogels and scatter plot graph (right) depicting migration speed (n=3, N > 20). See also Video S3, S4. (D) schematic depiction of PKM2 activation-driven phenotypic alterations. Error bars denote mean□±□SEM. Unpaired Student’s□*t*□test was performed for statistical significance (\**P*□≤ 0.05, \*\**P*□≤ 0.01, \*\*\**P*□< 0.001, \*\*\*\**P*□< 0.0001).

### PKM2 inhibition decreases cancer migration on stiff matrices

We next asked if the converse of the above observations were true – did inhibition of PKM2 in cancer cells on stiffer substrata phenocopy the traits of counterparts in softer environments? We cultivated MDA-MB-231 cells on 20 kPa gels in the presence of PKM2-IN-1 (compound 3k), a small molecule functional inhibitor of PKM2 (Ning et al., 2017). PKM2 inhibition significantly increased its nuclear localization on stiff hydrogels (Figure 3A for MDA-MB-231 and, S3A for HCC-1608; white signal represents DNA stained by DAPI and red represents PKM2 antibody signals). Nuclear/cytoplasmic ratio in PKM2 levels was significantly increased after PKM2-IN-1 treatment for both MDA-MB-231 and HCC1806 (the translocation is mildly confounded by lower overall levels of PKM2 because of an autoregulatory feedback of the protein (J. Liang et al., 2016); Figure 3A right for MDA-MB-231 and Figure S2A right for HCC1806; p < 0.0001 for both the cells with significance assessed using unpaired Student’s t-test). Intriguingly, PKM2 inhibition also resulted in an increase in mean cell area (Figure 3B with white signal represents DNA stained by DAPI and green represents F-actin stained by phalloidin with p < 0.0001 for total cell area); this could be an effect of the overall depletion, and an opposite increase in aspect ratio (p = 0.01 with significance assessed using unpaired Student’s t-test) in MDA-MB-231 on stiff matrices. None of these morphometric parameters were significantly changed in HCC1806 (Figure S3B with white signal represents DNA stained by DAPI and green represents F-actin stained by phalloidin). However, timelapse videography for cell motility showed that PKM2 inhibition reduced cell migration speeds significantly on stiff matrices (Figure 3C for MDA-MB-231 and S3C for HCC-1608; migration tracked for RFP-expressing MDA-MB-231 and GFP-expressing HCC1806 cells with individual tracks shown in distinct colors; video S5 for MDA-MB-231 control cells, video S6 for MDA-MB-231 cells treated with PKM2 inhibitor, video S13 for HCC1806 control cells and video S14 for HCC1806 cells treated with PKM2 inhibitor on 20 kPa hydrogels respectively). We also confirmed that inhibition of PKM2 elevated the intra- and extracellular levels of lactate in MDA-MB-231 cells which suggests an elevated glycolytic flux associated with a relatively increased nuclear localization (Figure 3D, p = 0.0014, p = 0.0004 extracellular and intracellular lactate, significance assessed using unpaired Students t-test). These observations suggest that PKM2 inhibition reverses the high migrational dynamics of cancer cells on stiff substrata, even though it showed a less dramatic effect on cell shape polarization (schematic depiction of trait correlations under PKM2 inhibition shown in Figure 3E).

**Figure 3.**
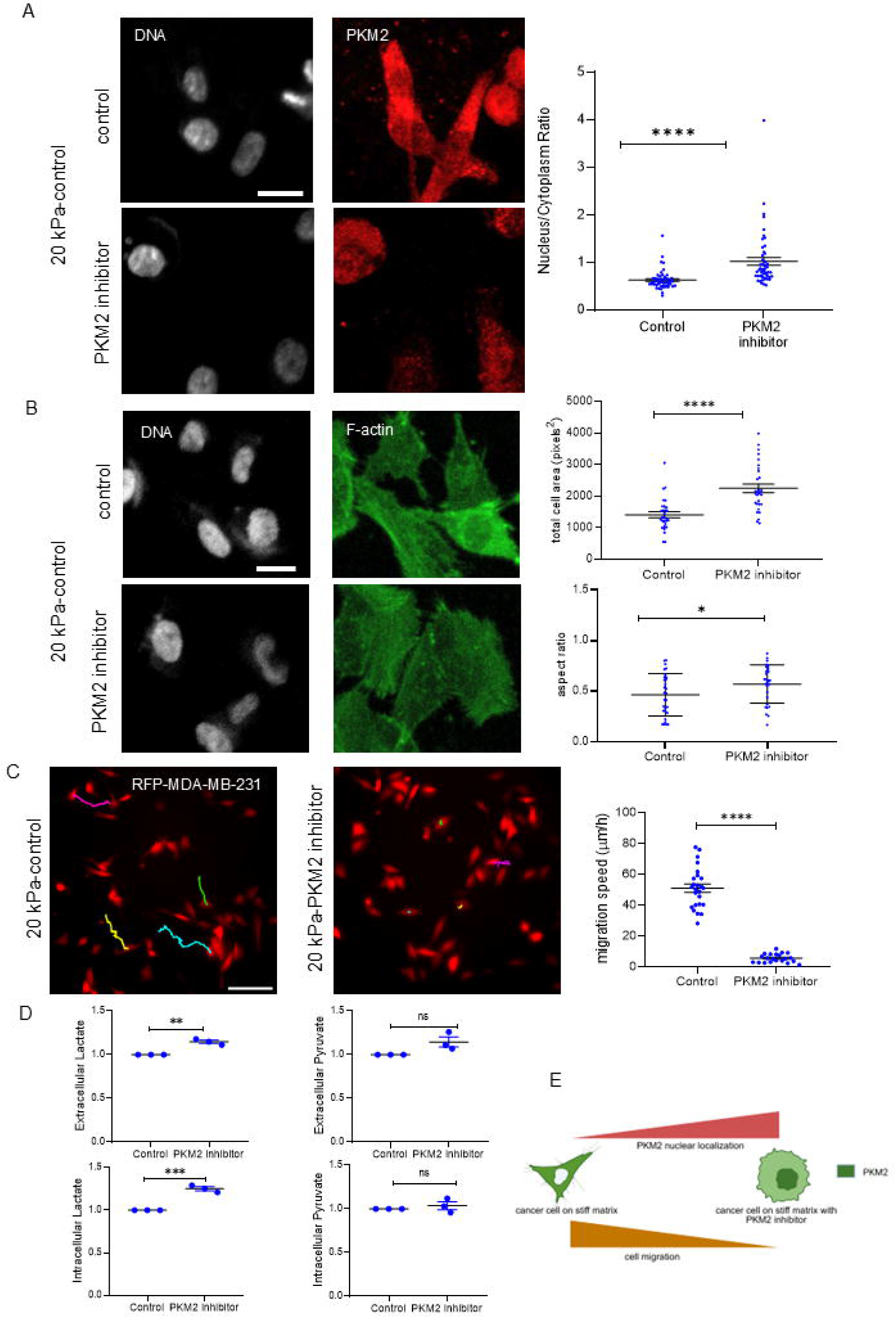
Decreased MDA-MB-231 breast cancer cell migration on stiff matrices upon PKM2 inhibition. (A) Representative confocal photomicrographs of breast cancer MDA-MB-231 cells untreated (top row of micrographs) or treated with 1.5LμM PKM2-IN-1, PKM2 inhibitor (second row from top) grown on 20 kPa (stiff) hydrogels coated with Collagen I and observed after 24□h of culture with staining by□DNA (white using DAPI, left panel) and PKM2 (Red, right panel). (n□= 3). Scale bar: 20□μm. Scatter plot graph (right) depicting nucleus/cytoplasm ratio of PKM2 between untreated and 1.5□μM PKM2-IN-1 treated MDA- MB-231 cells on 20 kPa hydrogels (n=3, N=30). (B) Representative confocal photomicrographs of breast cancer MDA-MB-231 cells untreated (top row of micrographs) or treated with 1.5□μM PKM2-IN-1 (second row from top) grown on 20 kPa (stiff) hydrogels coated with Collagen I and observed after 24□h of culture with staining by□DNA (white using DAPI, left panel) and F-actin (green using phalloidin, right panel (n□= 3). Scale bar: 20□μm. Graphs showing total cell area (right top) and aspect ratio (right bottom) of individual MDA-MB-231 cells untreated and treated with 1.5□μM PKM2-IN-1 on 20 kPa hydrogels (n□= 3, N = 30 single cells). (C) Epifluorescence photomicrographs of RFP- labelled MDA-MB-231 cells with their migration tracks for 3 hours, control (left) and treated with 1.5□μM PKM2-IN-1 (right) on 20 kPa hydrogels and scatter plot graph (right most) depicting migration speed (n=3, N > 20). See also Video S5, S6. (D) Graph measuring fold change of extracellular lactate (top left), extracellular pyruvate (top right), intracellular lactate (bottom left), intracellular pyruvate (bottom right) between untreated and 2□μM PKM2-IN-1 treated MDA-MB-231 cells on 20 kPa hydrogels (n=3). (E) Schematic depiction of alterations of traits under PKM2 inhibition. Error bars denote mean□±□SEM. Unpaired Student’s□*t*□test was performed for statistical significance (\**P*□≤ 0.05, \*\**P*□≤ 0.01, \*\*\**P*□< 0.001, \*\*\*\**P*□< 0.0001).

### PKM2 and YAP reciprocally regulate their subcellular repartitioning

Our observations suggested that the subcellular localization of PKM2 may act as a putative mechanosensing-mechanoresponsive signal for cancer cells allowing them to modulate their invasive traits based on the stiffness cues transduced from the external milieu. The Hippo pathway transcriptional regulator YAP has long been known to play such a role as well, and nuclear localization of YAP is observed in stiffer milieu (W. Chen et al., 2021; Ishihara & Haga, 2022). This led us to probe whether PKM2 and YAP localization are linked mechanistically in driving cellular response to rheological variation. We first confirmed higher nuclear localization of YAP in MDA-MB-231 and HCC1806 cells on stiffer hydrogels compared with soft counterparts (Figure 4A for MDA-MB-231 and S4A for HCC-1608; white signal represents DNA stained by DAPI and red represents YAP antibody signals; stiffness-cognate nucleocytoplasmic ratio of YAP shown in graph on the right for MDA-MB-231 in Figure 4A and in Figure S4A right for HCC1806 respectively, p = 0.0001 for MDA-MB-231, p = 0.0066 for HCC1806; significance assessed using unpaired Student’s t-test). To delineate whether PKM2 regulates YAP localization, we probed for YAP localization through immunocytochemistry on soft gels after treatment with the PKM2 cytoplasmic activator TEPP-46 and found greater nuclear localization compared with untreated controls (Figure 4B for MDA-MB-231 and S4B for HCC-1608; white signal represents DNA stained by DAPI and red represents YAP antibody signals; nuclear/cytoplasmic ratio of YAP assessed as a consequence of PKM2 activation in graph on the right for MDA-MB-231 with p < 0.0001 and Figure S4B for HCC1806 with p = 0.019; significance assessed using unpaired Student’s t-test). A schematic depiction of YAP localization as a function of PKM2 activation in soft gels is shown in Figure 4C.

**Figure 4.**
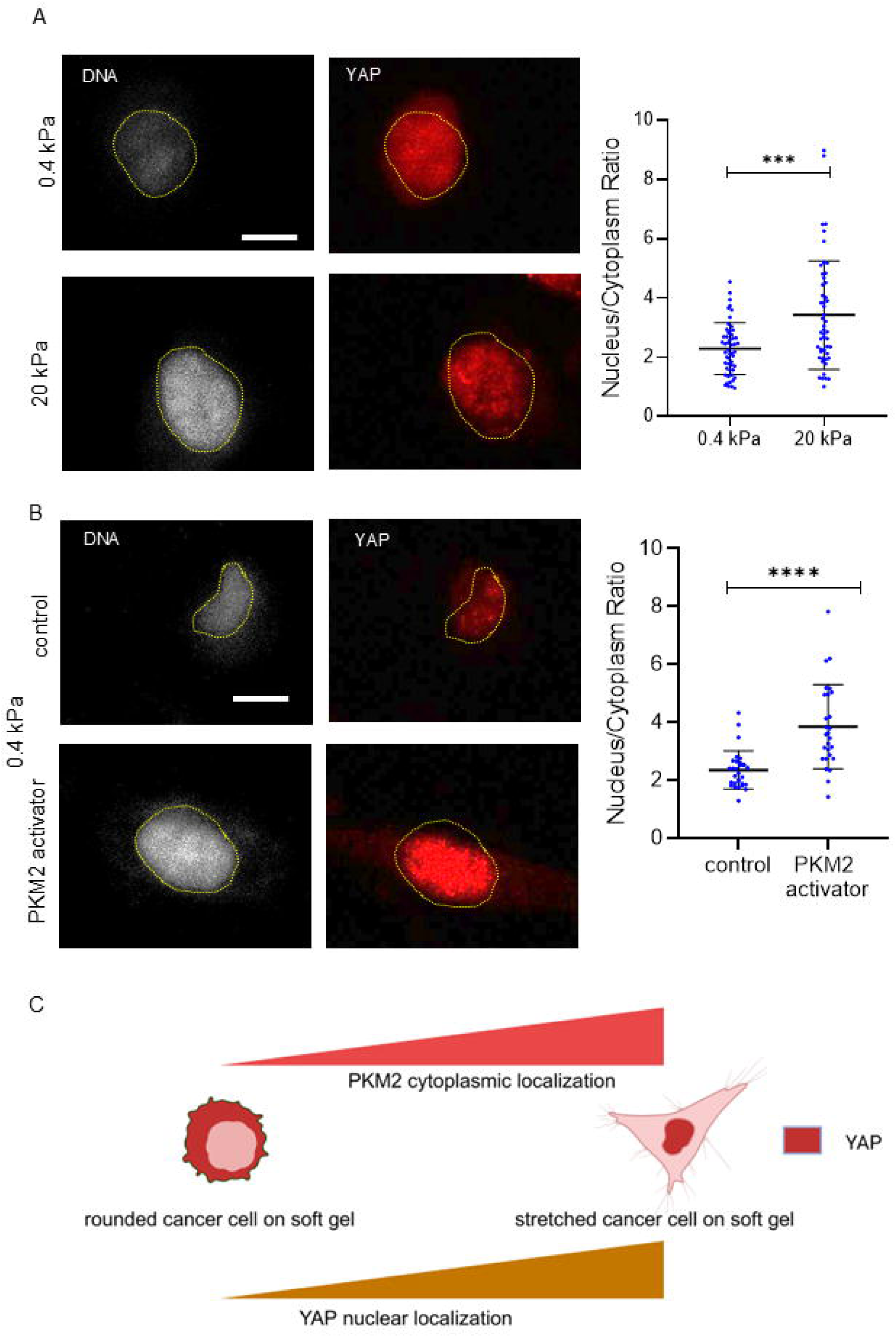
PKM2 regulates YAP localization on soft matrices in MDA-MB-231. (A) Representative confocal photomicrographs of breast cancer MDA-MB-231 cells grown on 0.4 kPa (top row of micrographs) and 20 kPa (bottom row of micrographs) polyacrylamide hydrogels coated with Collagen I and observed after 24□h of culture with staining for DNA (white using DAPI, left panel) and YAP (red, right panel). Yellow dotted areas represent nuclear area (n□= 3). Scale bar: 10□μm. Adjacent scatter plot graph depicting nucleus/cytoplasm ratio of YAP between MDA-MB-231 grown on 0.4 kPa and 20 kPa hydrogels (n = 3, n > 50). (B) Representative confocal photomicrographs of breast cancer MDA-MB-231 cells untreated (top row of micrographs) or treated with 40□μM TEPP-46, PKM2 cytoplasmic activator (second row from top) grown on 0.4 kPa (soft) hydrogels coated with Collagen I and observed after 24□h of culture with staining by□DNA (white using DAPI, left panel) and YAP (Red, right panel). Yellow dotted areas represent nuclear area (n□= 3). Scale bar: 10□μm. Adjacent scatter plot graph depicting nucleus/cytoplasm ratio of YAP between untreated and 40□μM TEPP-46 treated MDA-MB-231 cells on 0.4 kPa hydrogels (n = 3, N > 25). (C) Schematic depiction of YAP localization after PKM2 activation in soft gels. Error bars denote mean□±□SEM. Unpaired Student’s□*t*□test was performed for statistical significance (\**P*□≤ 0.05, \*\**P*□≤ 0.01, \*\*\**P*□< 0.001, \*\*\*\**P*□< 0.0001).

To further investigate if YAP regulates cell migration through PKM2 localization, we used verteporfin, an inhibitor of YAP signaling (Pan et al., 2016) and observed increased cellular circularity (Figure 5A for MDA-MB-231 and S5A for HCC-1608; white signal represents DNA stained by DAPI and green represents F-actin stained by phalloidin; p < 0.0001 for cell area for both the cells, p = 0.0012 for MDA-MB-231 and p < 0.0001 for HCC1806 respectively for aspect ratio; significance assessed using unpaired Students t-test), reduced total cell area and higher aspect axis ratio on stiff substrata indicating lower cell spreading and elongation. PKM2 localization after YAP inhibition showed increased nuclear localization and higher nuclear to cytoplasmic ratio was observed on stiff matrices (Figure 5B for MDA-MB-231 and S5B for HCC-1608, white signal represents DNA stained by DAPI and red represents cognate antibody signals; stiffness-cognate nucleocytoplasmic ratio of PKM2 shown in graph on the right, p < 0.0001; significance assessed using unpaired Student’s t-test). In addition, time lapse imaging revealed significant reduction in migration of MDA-MB-231 and HCC 1806 cells upon YAP inhibition (Figure 5C for MDA-MB-231 and S5C for HCC1806; migration tracked for RFP-expressing MDA-MB-231 and GFP-expressing HCC1806 cells with individual tracks shown in distinct colors; graph showing significantly higher mean migration speeds on 20 kPa; p < 0.0001; significance assessed using unpaired Students t-test; video S7 for MDA-MB-231 control cells, video S8 for MDA-MB-231 cells treated with YAP inhibitor, video S15 for HCC1806 control cells and video S16 for HCC1806 cells treated with YAP inhibitor on 20 kPa hydrogels respectively). Finally, lactate and pyruvate estimation assays also revealed that YAP inhibition increases both intracellular and extracellular lactate and pyruvate levels in MDA-MB-231 cells (Figure 5D; p = 0.0017, p < 0.0001, p < 0.0001 and p = 0.0076 for extracellular and intracellular lactate and pyruvate respectively, significance assessed using unpaired Students t-test) suggesting an increased glycolytic flux that phenocopies metabolic traits of cancer cells on soft matrices. These results not just suggest a reciprocal regulation in localization between PKM2 and YAP (Figure 5E), but also a modulation of the two molecules of each other’s canonical roles in mechanotransduction and metabolic flux regulation.

**Figure 5.**
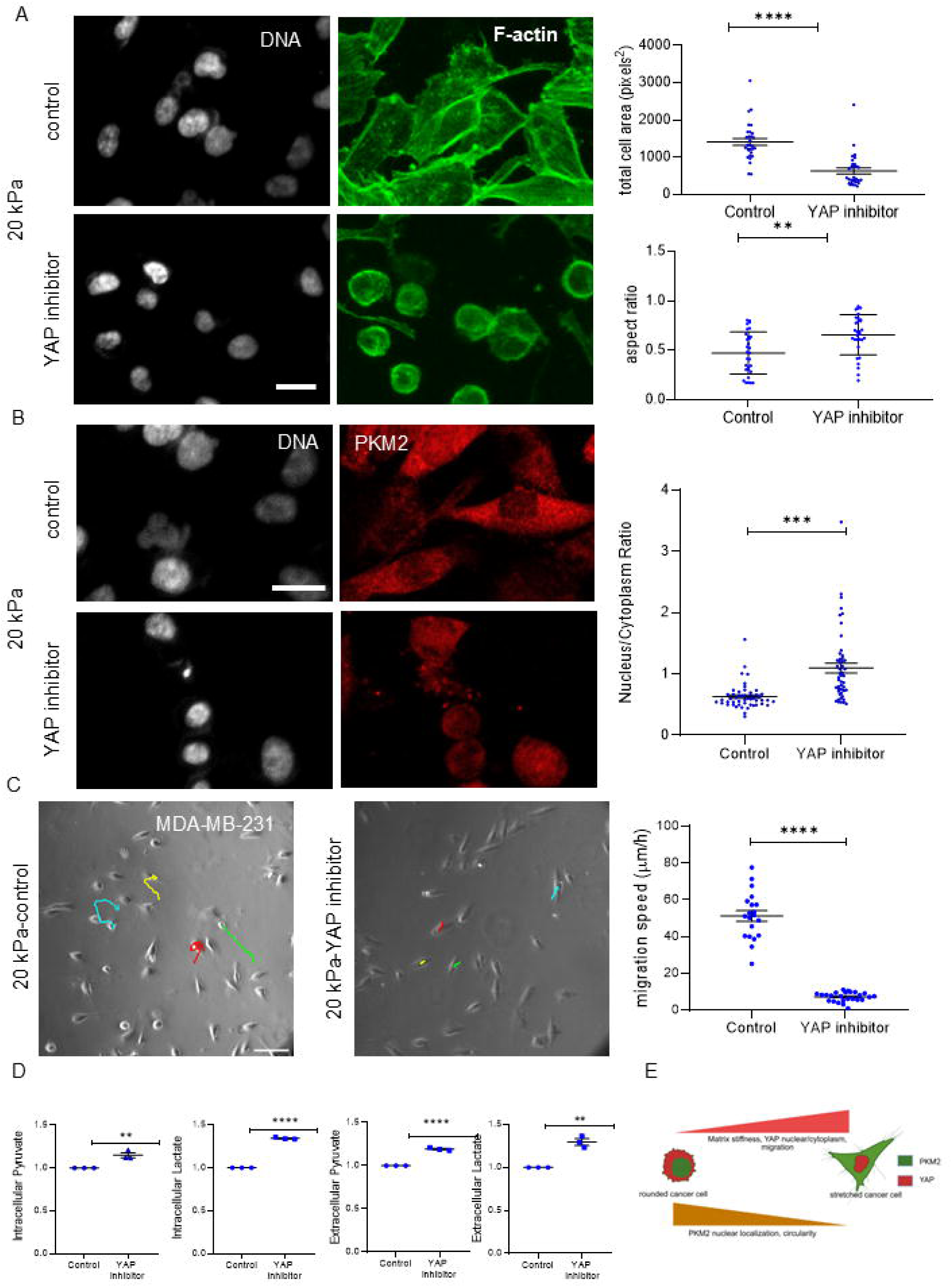
YAP regulates PKM2 localization on stiff matrices in MDA-MB-231. (A) Representative confocal photomicrographs of breast cancer MDA-MB-231 cells untreated (top row of micrographs) or treated with 5□μM Verteporfin, YAP signalling inhibitor (second row from top) grown on 20 kPa (stiff) hydrogels coated with Collagen I and observed after 24□h of culture with staining by□DNA (white using DAPI, left panel) and F-actin (green using phalloidin, right panel (n□= 3). Scale bar: 20□μm. Graphs showing total cell area (top) and aspect ratio (second from top) of individual MDA-MB-231 cells untreated and treated with 5□μM Verteporfin on 20 kPa hydrogels (n□= 3, N□= 30 single cells). (B) Representative confocal photomicrographs of MDA-MB-231 cells untreated (top row of micrographs) or treated with 5□μM Verteporfin (second row from top) grown on 20 kPa (stiff) hydrogels coated with Collagen I and observed after 24□h of culture with staining by□DNA (white using DAPI, left panel) and PKM2 (Red, right panel) and adjacent scatter plot graph (right) depicting nucleus/cytoplasm ratio of PKM2 between untreated and 5□μM Verteporfin treated MDA-MB-231 cells on 20 kPa hydrogels (n=3, N=50). (n□= 3). Scale bar: 20□μm. (C) Epifluorescence photomicrographs of MDA-MB-231 cells with their migration tracks for 3 hours, control (left) and treated with 5□μM Verteporfin (right) on 20 kPa hydrogels and scatter plot graph (right) depicting migration speed (n=3, N > 20). See also Video S7, S8. (D) Graph measuring fold change of intracellular pyruvate (first from left), intracellular lactate (second from left), extracellular pyruvate (third from left), extracellular lactate (last from left) between untreated and 5□μM Verteporfin treated MDA-MB-231 cells on 20 kPa hydrogels (n=3). (E) Schematic depiction of PKM2-YAP reciprocal repartitioning. Error bars denote mean□±□SEM. Unpaired Student’s□*t*□test was performed for statistical significance (\**P*□≤ 0.05, \*\**P*□≤ 0.01, \*\*\**P*□< 0.001, \*\*\*\**P*□< 0.0001).

### PKM2 and YAP inhibition decrease invasion of MDA-MB-231 spheroids in a 3D pathotypic multi-ECM microenvironment

The ability of PKM2 and YAP inhibitors to decrease breast cancer migration on stiff Collagen I substrata motivated to investigate whether inhibiton of these pathways also affect migration of cancer cells within three-dimensional Collagen I microenvironments. Clusters of MDA-MB-231 coated with laminin-rich basement membrane (lrBM) matrix and then embedded within Collagen I to mimic the were treated with 5 µM PKM2-IN-1 and 5 µM verteporfin (Pally, Banerjee, et al., 2022). Consistent with our single cell migration in 2D, compared to untreated controls (left) treatment with PKM2 inhibitor (middle) and with YAP inhibitor (right) were found to radially invade into the Type 1 collagen to a lower extent than control cancer cells (Figure 6A, brightfield images at 0 h, 8 h, 16 h and 24 h (top to bottom); video S17 for control cells, video S18 for cells treated with PKM2 inhibitor, video S19 for cells treated with YAP inhibitor). Time-lapse bright field microscopy on such cultures allowed us to measure the number of single cells that invaded into the collagen; normalized to initial spheroid size migrated cell number was found to be significantly lower after both PKM2 and YAP inhibition (Figure 6B, p = 0.0036 for control vs PKM2 inhibition and p = 0.0031 for control vs YAP inhibition respectively; significance assessed by one way ANOVA).

**Figure 6.**
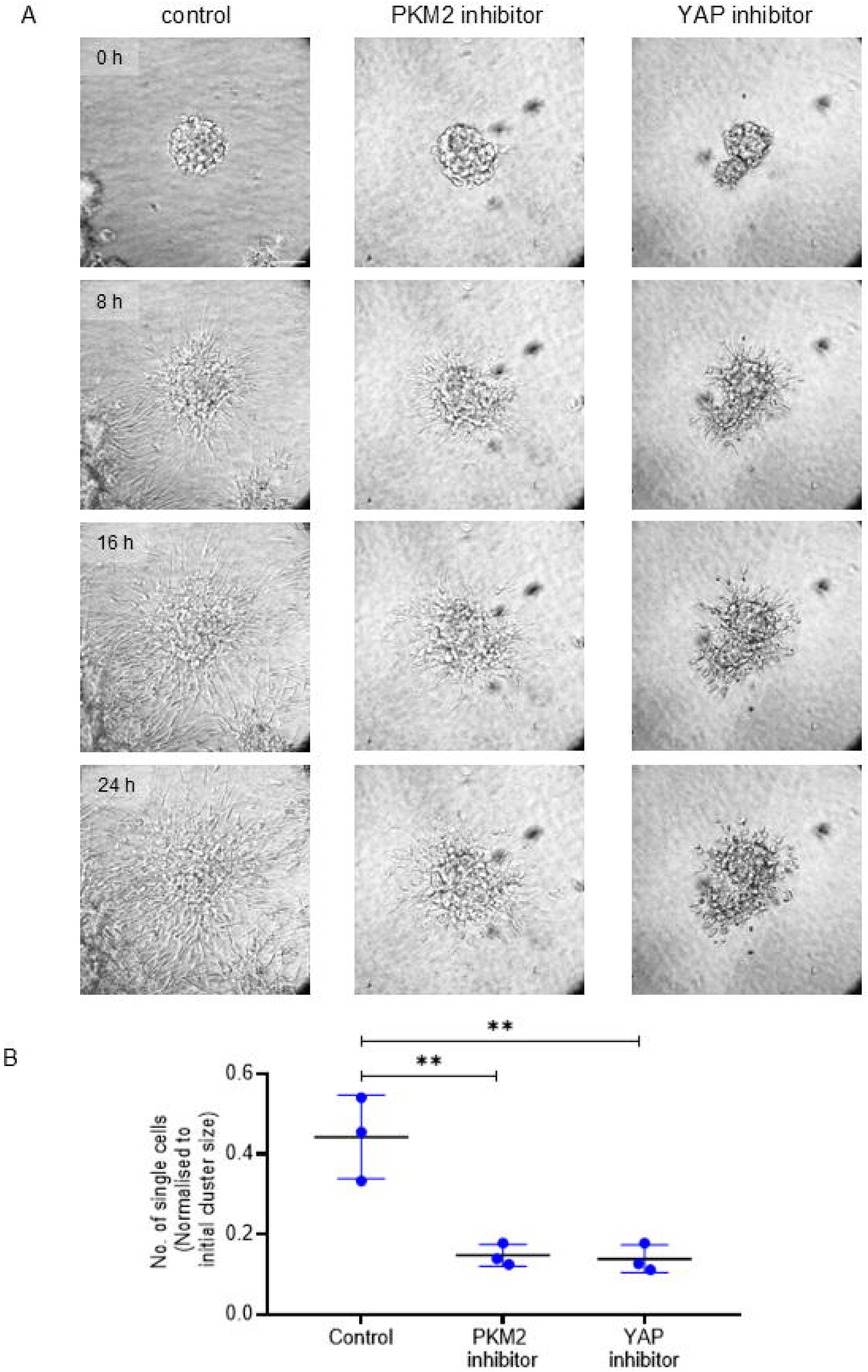
PKM2 and YAP inhibition decrease invasion of MDA-MB-231 spheroids through a 3D pathotypic multi-ECM microenvironment. (A) Bright field micrographs taken at 0, 8, 16, and 24 h (top to bottom) from time-lapse videography of lrECM-coated clusters of MDA-MB-231 cells invading into surrounding Collagen I, control (left), treated with 5 μM PKM2-IN-1(middle) and treated with 5 μM Verteporfin (right). (B) Scatter plot graph depicting number of dispersed single cells in Collagen I normalized to the initial cluster size obtained from time lapse videography (*n* = 3). Error bars denote mean ± SEM. One-way ANOVA with Tukey’s multiple comparison test was performed for statistical significance (\**P* ≤ 0.05, \*\**P* ≤ 0.01, \*\*\**P* < 0.001, \*\*\*\**P* < 0.0001). See also Video S17, S18 and S19.

## Discussion

In an insightful perspective on the links between metabolic and migration dynamics of cancer cells, Parlani and colleagues review how the bioenergetic stresses within tumor microenvironments as a consequence of nutrient deprivation, hypoxia and malperfusion coerce phenomenological transitions in motility strategies: from the consumption-heavy collective or mesenchymal modes to the relatively energy-favorable amoeboid behaviors (Parlani et al., 2022). Even without such dramatic transitions, normal and cancer cells have evolved more nuanced ways to co-locate sites of energy production and migration-based consumption, such as by AMPK or Miro-1-dependent mitochondrial movement to lamellipodial migration front as well as through tethering of glycolytic enzymes to F-actin (Bays et al., 2017; Schuler et al., 2017). These observations suggest a dependence on a cooperative flux of metabolites through glycolysis and oxidative phosphorylation to result in ATP production for mediating cell shape change, F-actin branching and actin treadmilling. Despite such advances, Parlani and colleagues pose as an outstanding question, how little is known about the joint regulation of the programs of energetic and migration dynamics so as to interlink the two on fast time scales. In this paper, we seek to shed insight on this question, through the novel demonstration of nucleocytoplasmic repartitioning of the rate limiting enzyme PKM2 as a response to the mechanical stiffness of the cancer cell substratum.

While ECM stiffness is known to modulate cancer migration, metastasis, and drug resistance (Acerbi et al., 2015; Ishihara & Haga, 2022; Li et al., 2023), its ability to regulate tumor metabolic dynamics is still ill understood. Here we provide direct evidence that matrix stiffness regulates the levels of glucose-uptake-derived metabolites in aggressive triple-negative breast cancer cells. Though cancer cells were once thought to rely on glycolysis under even aerobic conditions (Warburg’s effect), recent evidence shows that cancer cells deploy mitochondrial respiration in presence of glycolysis or shunt between glycolysis and mitochondrial oxidative phosphorylation in context dependent manner (Commander et al., 2020; DeBerardinis & Chandel, 2020; Warburg et al., 1927; J. Zhang et al., 2019). Such plasticity in metabolic strategies is essential for fulfilling both anabolic and energetic demands during cancer progression. However, how exactly such plasticity can be constrained by extraneous rheological cues are only beginning to be understood.

In the current study, consistent with previous results, we confirmed that matrix stiffening increases breast cancer motility and cell polarization. Park and co-workers have recently shown how human bronchial epithelial cells regulate a glycolytic rate-limiting enzyme phosphofructokinase (PFK) through actin cytoskeleton in a stiffness dependent manner (Park et al., 2020). Another report suggests a significant increase in aerobic glycolysis-regulating proteins with matrix stiffening in hepatocellular carcinoma cells (Liu et al., 2020). These observations prompted us to assay for levels of glycolytic intermediates, lactate and pyruvate as a function of stiffness in our setup. We found greater levels of extracellular and intracellular lactate and pyruvate on soft gels compared to stiff gels. This indicates that less migratory and deformable cells on soft gels prefer aerobic glycolysis which ends with lactate synthesis and secretion, whereas the more deformable and motile cancer cells preferentially funnel glycolytic intermediates towards oxidative phosphorylation and/or hexose monophosphate shunt. This is consistent with the reports of heightened lamellipodial spatialization of mitochondrial dynamics in migrative contexts (Madan et al., 2022; Zhao et al., 2013). In addition, we observe that PKM2, the final rate-limiting enzyme of glycolysis is predominantly localized in the nucleus on soft matrices whereas in stiffer substrata, it was present predominantly in the cytoplasm. PKM2 expression is known to be elevated in cancer cells, whereas the expression of other PKM isozymes PKM1, PK□, and PKR gradually decreases (Mazurek, 2008; Zahra et al., 2020). When present in cytoplasm, PKM2 exists as tetramer and catalyzes the synthesis of pyruvate at optimal levels, whereas, when it localizes to nucleus as dimer, it acts as a protein kinase or transcriptional co-activator to induce aerobic glycolysis and promote tumor progression. The low catalytic activity of dimeric PKM2 in nucleus results in increased production of glycolytic intermediates by inducing other glycolytic enzymes, the pentose phosphate pathway and glycerol synthesis and production of NADPH (Amin et al., 2019; Gao et al., 2012; Yang et al., 2011; Zahra et al., 2020). What’s more intriguing is that the modulation of PKM2 was observed to be downstream of, and transducing, rheological cues such that high PKM2 activity drove migration and shape polarization even on soft gels, where such traits in normal cells would be otherwise inhibited.

The regulation of migration by rheological alteration in ECM is canonically regulated by the Hippo pathway. A recent report indicates reversible regulation of YAP/TAZ pathway and glycolysis through PFK1 in regulating tumor progression (Enzo et al., 2015). Here we uncover, to the best of our knowledge for the first time, a mechanistic link between PKM2 and YAP localization in breast cancer cells in a stiffness-dependent manner: PKM2 cytoplasmic localization in soft gels increases YAP nuclear localization; the converse is true in stiffer milieu. Moreover, YAP signaling inhibition in stiff substrata not only reduces morphomigratory behavour of breast cancer cell but also increases PKM2 nuclear localization and both intracellular and extracellular lactate and pyruvate levels indicating an interlinking of mechanosensing and metabolic molecular machinery under rheological modulation.

It is pertinent to point out both the limitations of our study and how we intend to extend it in the future. Elastic modulus is not the only rheological trait of biological tissues and environments. Recent investigations not only suggest a viscoelastic nature to cellular ensembles, but the porosity of the ECM and interstitial fluid flow suggest poroelastic behavior operative at shorter time scales (Doyle et al., 2015; Salavati et al., 2022). Therefore, mechanical heterogeneities in tissues are a function of such rheological metrics, architectural features such as ECM porosity and biochemical variegation such as differential composition of ECM proteins (Ao et al., 2015; Provenzano et al., 2006; Wolf et al., 2013). Using more complex bioengineered setups with tunable trait variation will allow us to rigorously quantify the links between metabolic reprogramming and mechano-microenvironmental transduction. Secondly, what drives the repartitioning of PKM2 between nucleus and cytoplasm still remains elusive. That PKM2 does not have a canonical nuclear localization signal suggests it could be binding to other proteins that import it into the nucleus. What such proteins could be, and how they shuttle as a function of stiffness is a fruitful line of investigation for the future. In this study, we have unravelled the stiffness-dependent relocalization of only one enzyme, PKM2: it would be valuable to investigate the localization dynamics of other rate limiting enzymes of glycolysis, oxidative phosphorylation and hexose monophosphate shunt to decipher whether are our observations are part of a larger protein repartitioning that occurs to reorient metabolic flux based on mechanical context. These limitations notwithstanding, our results undeniably suggest mutual reinforcement of PKM2 and YAP in regulating the morphomigrational traits of breast cancer cell migration. Many YAP/TAZ signalling inhibitors are currently in clinical trials (Tolcher et al., 2022; Yap et al., 2023). Use of PKM2 inhibitors along with those YAP/TAZ inhibitors may hold the key to future therapeutic strategies. Our work proposes that interfering with cell mechanical sensing and metabolic dynamics may have broader implications in cancer therapy than previously recognized.

## Material and methods

### Cell culture and reagents

MDA-MB-231 cells were maintained in DMEM:F12 (1:1) (HiMedia, AT140) supplemented with 10% fetal bovine serum (FBS, 10270, Gibco). HCC1806 cells were cultured in RPMI-1640 medium (A□162A, HiMedia) along with 10% FBS (10270, Gibco). Both the cells were grown in a 37° C humidified incubator with 5% carbon dioxide. PKM2 cytoplasmic activator TEPP-46 (HY-18657, MedChemExpress), PKM2 inhibitor PKM2-IN-1 (HY-103617, MedChemExpress) and YAP inhibitor Verteporfin (HY-B0146, MedChemExpress) were used for the current study.

### Polyacrylamide (PA) gel preparation

PA gels of required stiffness were prepared in reference to the relative concentrations mentioned in a previous elegant study (Tse & Engler, 2010)

a. Activation of glass slide Silanization of glass slides (8-well chamber slides and 35 mm dishes with 20 mm glass bottom coverslips) was performed using 10% (3 Aminopropyl) trioethoxysilane (440140, Sigma) for 20 minutes followed by two washes with autoclaved filtered water. Fixation was then performed using 0.5% glutaraldehyde for 45 minutes and washed the same thrice with autoclaved filtered water.
b. Preparation of sandwich coverslip
c. top coverslip was coated with Rain-X to provide a hydrophobic coating followed by rinsing twice with autoclaved filtered water after 10 minutes.
d. Preparation of hydrogel(s) A solution of acrylamide and bis-acrylamide of the required concentration for specific stiffness was prepared in autoclaved filtered water. For 0.4 kPa gels, 3% acrylamide and 0.06% bis-acrylamide for 20 kPa gels, 8% acrylamide and 0.264% bis-acrylamide were used for this study. 60-80 μl of polymerizing gel solution per well for 8 chambered wells and 150 μl for 35 mm dishes with 20 mm glass coverslips is usually added. Therefore, to the required volume of acrylamide and bis-acrylamide solution, 1/100th of the total volume of 10% ammonium persulfate (APS) and 1/1000th of the total volume of tetramethylethylenediamine (TEMED) were added to produce polymerizing gel. This polymerizing gel was immediately added to the activated glass slide and the hydrophobic sandwich coverslip was placed immediately on top of it to spread the gel uniformly in the glass slide. Finally, the coverslip is removed after 30 minutes, as the gel usually forms by that time.
e. ECM coating of the gel surface The gel is initially coated with sulfo-SANPAH (803332, Sigma), a heterobifunctional photoreactive protein cross-linker for ECM coating. The coated gel is then exposed to UV for 15 mins followed by three phosphate buffer solution (PBS) washes to remove excess sulfo-SANPAH. To the activated gels, 100 μg/ml of neutralized collagen was added to 8 well chamber well or 35 mm glass bottom dishes and kept in the incubator at 37°C for 45 – 60 minutes for fast polymerization. Dishes were then transferred into the refrigerator at 4°C overnight to enhance the duration for attachment of collagen protein to the gel surface. The well-chambers are again placed in the incubator 1 hour prior to seeding cells for the experiment to equilibrate the temperature.

### Immunostaining

Cells were washed with 1X PBS after appropriate treatment and were fixed with 3.7 % of formaldehyde ((24005, Thermo Fisher Scientific)) for 20 minutes at 4°C. After fixation, cells were washed twice with 1X PBS and permeabilized using PBS with 0.5% Triton X-100 for 1 hour followed by blocking using PBS with 0.1% Triton X-100 and 3% BSA (MB083; HiMedia) for 45 min at RT. Primary antibody or Phalloidin incubation was carried out overnight in blocking buffer at 4°C. Subsequent processing was carried out in the dark. Following this, the cells were washed with PBS with 0.1 % Triton X-100 (PBST) thrice for 10 minutes each. Alexa Fluor^TM^ 568 (A-11011, Invitrogen) or Alexa Fluor^TM^ 488 (A-11008, Invitrogen) secondary antibody incubation was performed at room temperature under dark conditions for 2 hours. After PBST washes (10 min×3), cells were counterstained with DAPI (1:1000 dilution, D1306, Thermo Fisher Scientific) for 10 minutes. The cells are then washed with PBST thrice for 10 minutes each. Images were captured in 20× using a Carl Zeiss LSM880 laser confocal microscope. Images were processed and analyzed using Fiji image analysis software. Phalloidin-Alexa Fluor^TM^ 488 (1:500 dilution; A12379, Invitrogen) and Phalloidin-Alexa Fluor^TM^ 568 (1:500 dilution; A12380, Invitrogen) were used to stain F-actin. The primary antibodies used in our studies are against PKM2 (4053S, CST), YAP (14074S, CST).

### Timelapse experiment and tracking migration

Cells were grown on 8-well chambered cover glass with different stiffness matrices overnight. Following that, cells were treated with TEPP-46 or PKM2-IN-1 or Verteporfin for 24 hours. Time-lapse imaging of migratory cancer cells was then performed on Olympus IX83 Inverted Epi-fluorescence microscope fitted with stage top incubator and 5% carbon dioxide. Images were collected for 3h with every 3 min interval. Once the timelapse videos were obtained, single cell motility was tracked using MTrackJ plugin in Fiji.

### Lactate and pyruvate production assay

8×10^4^ cells were allowed to attach on 20 mm glass cover slip of 35 mm dishes of different stiffnesses overnight and inhibitors, activator treatment were carried out for 24 hours. After treatment, conditioned media and cell pellet were collected for extracellular and intracellular lactate (700510, Cayman) and pyruvate (700470, Cayman) assay respectively following the manufacturer’s instructions. Briefly culture medium and cell pellet were deproteinized with 0.5M metaphosphoric acid (MPA). After centrifugation, acid was neutralized with potassium carbonate. 20 µl of each sample was used for individual lactate and pyruvate assay according to manufacturer’s protocol. Fluorescence was measured using a microplate fluorometer equipped with a 540 nm excitation/590 nm emission filter set. Values are then normalized to cell numbers.

### 3D invasion assay

3D invasion assay was performed as described previously (Pally, Banerjee, et al., 2022). Briefly, MDA-MB-231 clusters were made using 30,000 cells per 200 µl of defined medium (Blaschke et al., 1994) supplemented with 4% rBM in a polyHEMA-coated (Sigma, P3932) 96-well plate. After 48 h, clusters were collected and embedded in polymerizing rat tail collagen in a chambered cover glass. 3D cultures were grown in a 37 °C humidified incubator with 5% carbon dioxide. Bright field time-lapse imaging of invading cancer clusters was performed on an Olympus IX83 fluorescence microscope fitted with a stage top incubator and 5% carbon dioxide. Images were collected for 24 h with every 30 min interval.

### Statistical analysis

All experiments were performed in at least duplicates and independently repeated thrice. All data are represented as mean ± SEM. Prism software (GraphPad Prism 8.0) was used for the generation of graphs and analysis. For statistical analysis, unpaired student’s *t*-test or one-way ANOVA with Tukey’s multiple comparison test was performed. Statistical significance is represented using *P*-value: * *P* ≤ 0.05, \*\**P* ≤ 0.01, \*\*\**P* ≤ 0.001, \*\*\*\**P* < 0.0001.Student’sC*t*Ctest was performed for statistical significance (\**P*C≤ 0.05, \*\**P*C≤ 0.01, \*\*\**P*C< 0.001, \*\*\*\**P*C< 0.0001).

## Acknowledgement

This work was supported by the Wellcome Trust/DBT India Alliance Fellowship/Grant [IA/I/17/2/503312] awarded to R.B. It was also supported by the Indo-French Centre for the Promotion of Advanced Research (69T08-2) to R.B. M.B. acknowledges support from DBT-JRF/SRF Fellowship. M.P. and S.M. acknowledge support from the Kishore Vaigyanik Protsahan Yojana Fellowship and the Prime Ministers Research Fellowship respectively.

## Data availability statement

The data that support the findings of this study are available from the corresponding author upon reasonable request.

## References

Acerbi, I., Cassereau, L., Dean, I., Shi, Q., Au, A., Park, C., Chen, Y. Y., Liphardt, J., Hwang, E. S., & Weaver, V. M. (2015). Human breast cancer invasion and aggression correlates with ECM stiffening and immune cell infiltration. Integrative Biology (United Kingdom), 7(10), 1120–1134. 10.1039/C5IB00040H,

Amin, S., Yang, P., & Li, Z. (2019). Pyruvate kinase M2: A multifarious enzyme in non-canonical localization to promote cancer progression. Biochimica et Biophysica Acta - Reviews on Cancer, 1871(2), 331–341. 10.1016/j.bbcan.2019.02.003

Anastasiou, D., Yu, Y., Israelsen, W. J., Jiang, J. K., Boxer, M. B., Hong, B. S., Tempel, W., Dimov, S., Shen, M., Jha, A., Yang, H., Mattaini, K. R., Metallo, C. M., Fiske, B. P., Courtney, K. D., Malstrom, S., Khan, T. M., Kung, C., Skoumbourdis, A. P., … Vander Heiden, M. G. (2012). Pyruvate kinase M2 activators promote tetramer formation and suppress tumorigenesis. Nature Chemical Biology, 8(10), 839–847. 10.1038/NCHEMBIO.1060,

Ao, M., Brewer, B. M., Yang, L., Franco Coronel, O. E., Hayward, S. W., Webb, D. J., & Li, D. (2015). Stretching fibroblasts remodels fibronectin and alters cancer cell migration. Scientific Reports, 5(1), 8334. 10.1038/SREP08334;TECHMETA=101,62;SUBJMETA=166,631,639,80,985;KWRD=BIOMEDICAL+ENGINEERING,CELL+BIOLOGY

Baker, E. L., Bonnecaze, R. T., & Zaman, M. H. (2009). Extracellular Matrix Stiffness and Architecture Govern Intracellular Rheology in Cancer. Biophysical Journal, 97(4), 1013. 10.1016/J.BPJ.2009.05.054

Bays, J. L., Campbell, H. K., Heidema, C., Sebbagh, M., & Demali, K. A. (2017). Linking E-cadherin mechanotransduction to cell metabolism through force-mediated activation of AMPK. Nature Cell Biology 2017 19:6, 19(6), 724–731. 10.1038/ncb3537

Blaschke, R. J., Howlett, A. R., Desprez, P. Y., Petersen, O. W., & Bissell, M. J. (1994). [25] Cell differentiation by extracellular matrix components. Methods in Enzymology, 245(C), 535–556. 10.1016/0076-6879(94)45027-7

Chandrashekar, D. S., Karthikeyan, S. K., Korla, P. K., Patel, H., Shovon, A. R., Athar, M., Netto, G. J., Qin, Z. S., Kumar, S., Manne, U., Crieghton, C. J., & Varambally, S. (2022). UALCAN: An update to the integrated cancer data analysis platform. Neoplasia (United States), 25, 18–27. 10.1016/j.neo.2022.01.001

Chen, W., Park, S., Patel, C., Bai, Y., Henary, K., Raha, A., Mohammadi, S., You, L., & Geng, F. (2021). The migration of metastatic breast cancer cells is regulated by matrix stiffness via YAP signalling. Heliyon, 7(2), e06252. 10.1016/J.HELIYON.2021.E06252

Chen, X., Chen, S., & Yu, D. (2020). Protein kinase function of pyruvate kinase M2 and cancer. Cancer Cell International, 20(1), 523. 10.1186/S12935-020-01612-1

Christofk, H. R., Vander Heiden, M. G., Harris, M. H., Ramanathan, A., Gerszten, R. E., Wei, R., Fleming, M. D., Schreiber, S. L., & Cantley, L. C. (2008). The M2 splice isoform of pyruvate kinase is important for cancer metabolism and tumour growth. Nature, 452(7184), 230–233. 10.1038/NATURE06734;KWRD=SCIENCE

Commander, R., Wei, C., Sharma, A., Mouw, J. K., Burton, L. J., Summerbell, E., Mahboubi, D., Peterson, R. J., Konen, J., Zhou, W., Du, Y., Fu, H., Shanmugam, M., & Marcus, A. I. (2020). Subpopulation targeting of pyruvate dehydrogenase and GLUT1 decouples metabolic heterogeneity during collective cancer cell invasion. Nature Communications 2020 11:1, 11(1), 1–17. 10.1038/s41467-020-15219-7

DeBerardinis, R. J., & Chandel, N. S. (2020). We need to talk about the Warburg effect. Nature Metabolism 2020 2:2, 2(2), 127–129. 10.1038/s42255-020-0172-2

Doostmohammadi, A., & Ladoux, B. (2022). Physics of liquid crystals in cell biology. Trends in Cell Biology, 32(2), 140–150. 10.1016/J.TCB.2021.09.012

Doyle, A. D., Carvajal, N., Jin, A., Matsumoto, K., & Yamada, K. M. (2015). Local 3D matrix microenvironment regulates cell migration through spatiotemporal dynamics of contractility-dependent adhesions. Nature Communications 2015 6:1, 6(1), 1–15. 10.1038/ncomms9720

Enzo, E., Santinon, G., Pocaterra, A., Aragona, M., Bresolin, S., Forcato, M., Grifoni, D., Pession, A., Zanconato, F., Guzzo, G., Bicciato, S., & Dupont, S. (2015). Aerobic glycolysis tunes YAP/TAZ transcriptional activity. The EMBO Journal, 34(10), 1349. 10.15252/EMBJ.201490379

Evans, K. W., Yuca, E., Scott, S. S., Zhao, M., Arango, N. P., Cruz Pico, C. X., Saridogan, T., Shariati, M., Class, C. A., Bristow, C. A., Vellano, C. P., Zheng, X., Gonzalez-Angulo, A. M., Su, X., Tapia, C., Chen, K., Akcakanat, A., Lim, B., Tripathy, D., … Meric-Bernstam, F. (2021). Oxidative Phosphorylation Is a Metabolic Vulnerability in Chemotherapy-Resistant Triple-Negative Breast Cancer. Cancer Research, 81(21), 5572. 10.1158/0008-5472.CAN-20-3242

Gao, X., Wang, H., Yang, J. J., Liu, X., & Liu, Z. R. (2012). Pyruvate Kinase M2 Regulates Gene Transcription by Acting as a Protein Kinase. Molecular Cell, 45(5), 598–609. 10.1016/j.molcel.2012.01.001

Göransson, S., Olofsson, H., Johansson, H. J., Yan, F., Vogiatzakis, C., Liang, S., Bellato, H. M., Masvidal, L., Aksoylu, I., Hartman, J., Hajj, G. N., Larsson, O., Lehtiö, J., & Strömblad, S. (2025). Mechanical control of breast cancer malignancy by promotion of mevalonate pathway enzyme synthesis. Matrix Biology, 140, 1–15. 10.1016/j.matbio.2025.05.005

Hensley, C. T., Faubert, B., Yuan, Q., Lev-Cohain, N., Jin, E., Kim, J., Jiang, L., Ko, B., Skelton, R., Loudat, L., Wodzak, M., Klimko, C., McMillan, E., Butt, Y., Ni, M., Oliver, D., Torrealba, J., Malloy, C. R., Kernstine, K., … DeBerardinis, R. J. (2016). Metabolic Heterogeneity in Human Lung Tumors. Cell, 164(4), 681–694. 10.1016/j.cell.2015.12.034

Ishihara, S., & Haga, H. (2022). Matrix Stiffness Contributes to Cancer Progression by Regulating Transcription Factors. Cancers, 14(4). 10.3390/CANCERS14041049,

Jiang, J. kang, Boxer, M. B., Vander Heiden, M. G., Shen, M., Skoumbourdis, A. P., Southall, N., Veith, H., Leister, W., Austin, C. P., Park, H. W., Inglese, J., Cantley, L. C., Auld, D. S., & Thomas, C. J. (2010). Evaluation of Thieno[3,2-b]pyrrole[3,2-d]pyridazinones as Activators of the Tumor Cell Specific M2 Isoform of Pyruvate Kinase. Bioorganic & Medicinal Chemistry Letters, 20(11), 3387. 10.1016/J.BMCL.2010.04.015

Li, T., Li, Y., Wu, H., Peng, C., Wang, J., Chen, S., Zhao, T., Li, S., Qin, X., & Liu, Y. (2023). Extracellular cell matrix stiffness-driven drug resistance of breast cancer cells via EGFR activation. Mechanobiology in Medicine, 1(2), 100023. 10.1016/J.MBM.2023.100023

Liang, J., Cao, R., Zhang, Y., Xia, Y., Zheng, Y., Li, X., Wang, L., Yang, W., & Lu, Z. (2016). PKM2 dephosphorylation by Cdc25A promotes the Warburg effect and tumorigenesis. Nature Communications, 7, 12431. 10.1038/NCOMMS12431

Liang, L. J., Yang, F. Y., Wang, D., Zhang, Y. F., Yu, H., Wang, Z., Sun, B. B., Liu, Y. T., Wang, G. Z., & Zhou, G. B. (2024). CIP2A induces PKM2 tetramer formation and oxidative phosphorylation in non-small cell lung cancer. Cell Discovery, 10(1), 1–22. 10.1038/S41421-023-00633-0;SUBJMETA=2327,304,631,67,80;KWRD=CANCER+METABOLISM,MECHANISMS+OF+DISEASE

Liu, Q. P., Luo, Q., Deng, B., Ju, Y., & Song, G. Bin. (2020). Stiffer Matrix Accelerates Migration of Hepatocellular Carcinoma Cells through Enhanced Aerobic Glycolysis Via the MAPK-YAP Signaling. Cancers 2020, Vol. 12, Page 490, 12(2), 490. 10.3390/CANCERS12020490

Madan, S., Uttekar, B., Chowdhary, S., & Rikhy, R. (2022). Mitochondria Lead the Way: Mitochondrial Dynamics and Function in Cellular Movements in Development and Disease. Frontiers in Cell and Developmental Biology, 9. 10.3389/FCELL.2021.781933,

Mazurek, S. (2008). Pyruvate Kinase Type M2: A Key Regulator Within the Tumour Metabolome and a Tool for Metabolic Profiling of Tumours. Ernst Schering Foundation Symposium Proceedings, 4, 99–124. 10.1007/2789_2008_091

Mierke, C. T. (2021). Viscoelasticity Acts as a Marker for Tumor Extracellular Matrix Characteristics. Frontiers in Cell and Developmental Biology, 9, 785138. 10.3389/FCELL.2021.785138/XML/NLM

Morris, B. A., Burkel, B., Ponik, S. M., Fan, J., Condeelis, J. S., Aguire-Ghiso, J. A., Castracane, J., Denu, J. M., & Keely, P. J. (2016). Collagen Matrix Density Drives the Metabolic Shift in Breast Cancer Cells. EBioMedicine, 13, 146–156. 10.1016/J.EBIOM.2016.10.012

Ning, X., Qi, H., Li, R., Li, Y., Jin, Y., McNutt, M. A., Liu, J., & Yin, Y. (2017). Discovery of novel naphthoquinone derivatives as inhibitors of the tumor cell specific M2 isoform of pyruvate kinase. European Journal of Medicinal Chemistry, 138, 343–352. 10.1016/j.ejmech.2017.06.064

Olson, M. F., Sahai, A. E., Olson, M. F., & Sahai, E. (2008). The actin cytoskeleton in cancer cell motility. Clinical & Experimental Metastasis 2008 26:4, 26(4), 273–287. 10.1007/S10585-008-9174-2

Pally, D., Banerjee, M., Hussain, S., Kumar, R. V., Petersson, A., Rosendal, E., Gunnarsson, L., Peterson, K., Leffler, H., Nilsson, U. J., & Bhat, R. (2022). Galectin-9 Signaling Drives Breast Cancer Invasion through Extracellular Matrix. ACS Chemical Biology, 17(6), 1376–1386. 10.1021/ACSCHEMBIO.1C00902/SUPPL_FILE/CB1C00902_SI_001.XLSX

Pally, D., Goutham, S., & Bhat, R. (2022). Extracellular matrix as a driver for intratumoral heterogeneity. Physical Biology, 19(4). 10.1088/1478-3975/AC6EB0,

Pan, W., Wang, Q., Zhang, Y., Zhang, N., Qin, J., Li, W., Wang, J., Wu, F., Cao, L., & Xu, G. (2016). Verteporfin can reverse the paclitaxel resistance induced by YAP over-expression in HCT-8/T cells without photoactivation through inhibiting YAP expression. Cellular Physiology and Biochemistry, 39(2), 481–490. 10.1159/000445640,

Park, J. S., Burckhardt, C. J., Lazcano, R., Solis, L. M., Isogai, T., Li, L., Chen, C. S., Gao, B., Minna, J. D., Bachoo, R., DeBerardinis, R. J., & Danuser, G. (2020). Mechanical regulation of glycolysis via cytoskeleton architecture. Nature, 578(7796), 621. 10.1038/S41586-020-1998-1

Parlani, M., Jorgez, C., & Friedl, P. (2022). Plasticity of cancer invasion and energy metabolism. Trends in Cell Biology, 33(5), 388. 10.1016/J.TCB.2022.09.009

Provenzano, P. P., Eliceiri, K. W., Campbell, J. M., Inman, D. R., White, J. G., & Keely, P. J. (2006). Collagen reorganization at the tumor-stromal interface facilitates local invasion. BMC Medicine, 4(1). 10.1186/1741-7015-4-38,

Salavati, H., Debbaut, C., Pullens, P., & Ceelen, W. (2022). Interstitial fluid pressure as an emerging biomarker in solid tumors. Biochimica et Biophysica Acta (BBA) - Reviews on Cancer, 1877(5), 188792. 10.1016/J.BBCAN.2022.188792

Schuler, M. H., Lewandowska, A., Di Caprio, G., Skillern, W., Upadhyayula, S., Kirchhausen, T., Shaw, J. M., & Cunniff, B. (2017). Miro1-mediated mitochondrial positioning shapes intracellular energy gradients required for cell migration. Molecular Biology of the Cell, 28(16), 2159–2169. 10.1091/MBC.E16-10-0741/ASSET/IMAGES/LARGE/2159FIG6.JPEG

Sleeboom, J. J. F., van Tienderen, G. S., Schenke-Layland, K., van der Laan, L. J. W., Khalil, A. A., & Verstegen, M. M. A. (2024). The extracellular matrix as hallmark of cancer and metastasis: From biomechanics to therapeutic targets. Science Translational Medicine, 16(728), 3840. 10.1126/SCITRANSLMED.ADG3840,

So, W. Y., & Tanner, K. (2021). Emerging principles of cancer biophysics. Faculty Reviews, 10. 10.12703/R/10-61

Tien, J., Ghani, U., Dance, Y. W., Seibel, A. J., Karakan, M. Ç., Ekinci, K. L., & Nelson, C. M. (2020). Matrix Pore Size Governs Escape of Human Breast Cancer Cells from a Microtumor to an Empty Cavity. IScience, 23(11). 10.1016/j.isci.2020.101673

Tolcher, A. W., Lakhani, N. J., McKean, M., Lingaraj, T., Victor, L., Sanchez-Martin, M., Kacena, K., Malek, K. S., & Santillana, S. (2022). A phase 1, first-in-human study of IK-930, an oral TEAD inhibitor targeting the Hippo pathway in subjects with advanced solid tumors. Journal of Clinical Oncology, 40(16_suppl), TPS3168–TPS3168. 10.1200/JCO.2022.40.16_SUPPL.TPS3168

Tse, J. R., & Engler, A. J. (2010). Preparation of hydrogel substrates with tunable mechanical properties. Current Protocols in Cell Biology, Chapter 10(SUPPL. 47). 10.1002/0471143030.CB1016S47,

Uslu, C., Kapan, E., & Lyakhovich, A. (2025). OXPHOS inhibition overcomes chemoresistance in triple negative breast cancer. Redox Biology, 83, 103637. 10.1016/J.REDOX.2025.103637

Wang, T. C., Abolghasemzade, S., McKee, B. P., Singh, I., Pendyala, K., Mohajeri, M., Patel, H., Shaji, A., Kersey, A. L., Harsh, K., Kaur, S., Dollahon, C. R., Chukkapalli, S., Lele, P. P., Conway, D. E., Gaharwar, A. K., Dickinson, R. B., & Lele, T. P. (2024). Matrix stiffness drives drop like nuclear deformation and lamin A/C tension-dependent YAP nuclear localization. Nature Communications 2024 15:1, 15(1), 1–17. 10.1038/s41467-024-54577-4

Warburg, O., Wind, F., & Negelein, E. (1927). The metabolism of tumors in the body. Journal of General Physiology, 8(6), 519–530. 10.1085/JGP.8.6.519,

Winkler, J., Abisoye-Ogunniyan, A., Metcalf, K. J., & Werb, Z. (2020). Concepts of extracellular matrix remodelling in tumour progression and metastasis. Nature Communications, 11(1). 10.1038/S41467-020-18794-X,

Wolf, K., te Lindert, M., Krause, M., Alexander, S., te Riet, J., Willis, A. L., Hoffman, R. M., Figdor, C. G., Weiss, S. J., & Friedl, P. (2013). Physical limits of cell migration: Control by ECM space and nuclear deformation and tuning by proteolysis and traction force. Journal of Cell Biology, 201(7), 1069–1084. 10.1083/JCB.201210152,

Wozniak, M. A., Desai, R., Solski, P. A., Der, C. J., & Keely, P. J. (2003). ROCK-generated contractility regulates breast epithelial cell differentiation in response to the physical properties of a three-dimensional collagen matrix. Journal of Cell Biology, 163(3), 583–595. 10.1083/JCB.200305010,

Xu, R., Yin, P., Wei, J., & Ding, Q. (2023). The role of matrix stiffness in breast cancer progression: a review. Frontiers in Oncology, 13, 1284926. 10.3389/FONC.2023.1284926

Yang, W., Xia, Y., Ji, H., Zheng, Y., Liang, J., Huang, W., Gao, X., Aldape, K., & Lu, Z. (2011). Nuclear PKM2 regulates β-catenin transactivation upon EGFR activation. Nature, 480(7375), 118–122. 10.1038/NATURE10598,

Yap, T. A., Kwiatkowski, D. J., Desai, J., Dagogo-Jack, I., Millward, M., Kindler, H. L., Tolcher, A. W., Frentzas, S., Thurston, A. W., Post, L., & Dorr, F. A. (2023). Abstract CT006: First-in-class, first-in-human phase 1 trial of VT3989, an inhibitor of yes-associated protein (YAP)/transcriptional enhancer activator domain (TEAD), in patients (pts) with advanced solid tumors enriched for malignant mesothelioma and other tumors with neurofibromatosis 2 (NF2) mutations. Cancer Research, 83(8_Supplement), CT006–CT006. 10.1158/1538-7445.AM2023-CT006

Zahra, K., Dey, T., Ashish, Mishra, S. P., & Pandey, U. (2020). Pyruvate Kinase M2 and Cancer: The Role of PKM2 in Promoting Tumorigenesis. Frontiers in Oncology, 10, 505842. 10.3389/FONC.2020.00159/BIBTEX

Zhang, J., Goliwas, K. F., Wang, W., Taufalele, P. V., Bordeleau, F., & Reinhart-King, C. A. (2019). Energetic regulation of coordinated leader–follower dynamics during collective invasion of breast cancer cells. Proceedings of the National Academy of Sciences of the United States of America, 116(16), 7867–7872. 10.1073/PNAS.1809964116/SUPPL_FILE/PNAS.1809964116.SM03.MOV

Zhang, Z., Deng, X., Liu, Y., Liu, Y., Sun, L., & Chen, F. (2019). PKM2, function and expression and regulation. Cell & Bioscience 2019 9:1, 9(1), 1–25. 10.1186/S13578-019-0317-8

Zhao, J., Zhang, J., Yu, M., Xie, Y., Huang, Y., Wolff, D. W., Abel, P. W., & Tu, Y. (2013). Mitochondrial dynamics regulates migration and invasion of breast cancer cells. Oncogene, 32(40), 4814–4824. 10.1038/ONC.2012.494,

