## Supplementary figure legends for "A PKM2-YAP reciprocal repartitioning modulates invasion of breast cancer cells"

**Figure S1.** Matrix stiffness regulates morpho-migration dynamics and PKM2 localization of HCC1806 breast cancer cells. (A) Figure depicting analysis of nucleus/cytoplasm ratio. Yellow dotted lines indicate nuclear are and white dotted lines indicate total cell area.(B) Representative confocal photomicrographs of breast cancer HCC1806 cells grown on 0.4 kPa (top row of micrographs) and 20 kPa (bottom row of micrographs) polyacrylamide hydrogels coated with Collagen I and observed after 24 h of culture with staining for DNA (white using DAPI, left panel) and F-actin (green using phalloidin, right panel (n = 3). Scale bar: 20 μm. Adjacent graphs showing cell area (top) and aspect ratio (second from top) of individual HCC1806 cells grown on 0.4 kPa and 20 kPa hydrogels (n = 3, N = 30 single cells). (C) Epifluorescence photomicrographs of GFP labelled HCC1806 cells with their migration tracks for 3 hours, 0.4 kPa (left) and 20 kPa (right) hydrogels and scatter plot graph depicting migration speed (n=3, N > 20). See also Video S9, S10. (D) UALCAN TCGA sample analysis between normal breast and breast tumor for PKM2 expression. (E) Representative confocal photomicrographs of HCC18006 cells grown on 0.4 kPa (top row of micrographs) and 20 kPa (bottom row of micrographs) polyacrylamide hydrogels coated with Collagen I and observed after 24 h of culture with staining for DNA (white using DAPI, left panel) and PKM2 (Red, right panel). (n = 3). Scale bar: 20 μm and scatter plot graph (right) depicting nucleus/cytoplasm ratio of PKM2 (n=3, N=30). Error bars denote mean ± SEM. The unpaired Student’s *t* test was performed for statistical significance (**P* ≤ 0.05, ***P* ≤ 0.01, ****P* < 0.001, *****P* < 0.0001).

**Figure S2.** Increased HCC1806 breast cancer cell migration on soft matrices upon PKM2 cytoplasmic activation. (A) Representative confocal photomicrographs of breast HCC1806 cells untreated (top row of micrographs) or treated with 40 μM TEPP-46, PKM2 cytoplasmic activator (second row from top) grown on 0.4 kPa (soft) hydrogels coated with collagen I and observed after 24 h of culture with staining by DNA (white using DAPI, left panel) and PKM2 (Red, right panel), (n = 3). Scale bar: 20 μm. Scatter plot graph depicting nucleus/cytoplasm ratio of PKM2 between untreated and 40 μM TEPP-46 treated HCC1806 cells on 0.4 kPa hydrogels (n=3, N=30). (B) Representative confocal photomicrographs of breast cancer HCC1806 cells untreated (top row of micrographs) or treated with 40 μM TEPP-46, PKM2 cytoplasmic activator (second row from top) grown on 0.4 kPa (soft) hydrogels coated with collagen I and observed after 24 h of culture with staining by DNA (white using DAPI, left panel) and F-actin (green using phalloidin, right panel (n = 3). Scale bar: 20 μm. Adjacent graphs showing cell area (top right) and aspect ratio (bottom) of individual HCC1806 cells untreated and treated with 40 μM TEPP-46 on 0.4 kPa hydrogels (n = 3, N = 30 single cells). (C) Epifluorescence photomicrographs of RFP labelled HCC1806 cells with their migration tracks for 3 hours, control (left) and treated with 40 μM TEPP-46 (right) on 0.4 kPa hydrogels and scatter plot graph depicting migration speed (n=3, N > 20). See also Video S11, S12. Error bars denote mean ± SEM. The unpaired Student’s *t* test was performed for statistical significance (**P* ≤ 0.05, ***P* ≤ 0.01, ****P* < 0.001, *****P* < 0.0001).

**Figure S3.** Decreased HCC1806 breast cancer cell migration on stiff matrices upon PKM2 inhibition. (A) Representative confocal photomicrographs of breast cancer HCC1806 cells untreated (top row of micrographs) or treated with 0.5 μM PKM2-IN-1, PKM2 inhibitor (second row from top) grown on 20 kPa (stiff) hydrogels coated with collagen I and observed after 24 h of culture with staining by DNA (white using DAPI, left panel) and PKM2 (Red, right panel), (n = 3). Scale bar: 20 μm. Scatter plot graph depicting nucleus/cytoplasm ratio of PKM2 between untreated and 0.5 μM PKM2-IN-1 treated HCC1806 cells on 20 kPa hydrogels (n=3, N>20). (B) Representative confocal photomicrographs of HCC1806 cells untreated (top row of micrographs) or treated with 0.5 μM PKM2-IN-1 (second row from top) grown on 20 kPa (stiff) hydrogels coated with collagen I and observed after 24 h of culture with staining by DNA (white using DAPI, left panel) and F-actin (green using phalloidin, right panel (n = 3). Scale bar: 20 μm. Adjacent graphs showing cell area (top) and aspect ratio (second from top) of individual HCC1806 cells untreated and treated with 0.5 μM PKM2-IN-1 on 20 kPa hydrogels (n = 3, N > 20 single cells). (C) Epifluorescence photomicrographs of GFP labelled HCC1806 cells with their migration tracks for 3 hours, control (left) and treated with 0.5 μM PKM2-IN-1 (right) on 20 kPa hydrogels and scatter plot graph depicting migration speed (n=3, N > 20). See also Video S13, S14. Error bars denote mean ± SEM. The unpaired Student’s *t* test was performed for statistical significance (**P* ≤ 0.05, ***P* ≤ 0.01, ****P* < 0.001, *****P* < 0.0001).

**Figure S4**. PKM2 regulates YAP localization on soft matrices in HCC1806. (A) Representative confocal photomicrographs of breast cancer HCC1806 cells grown on 0.4 kPa (top row of micrographs) and 20 kPa (bottom row of micrographs) polyacrylamide hydrogels coated with Collagen I and observed after 24 h of culture with staining for DNA (white using DAPI, left panel) and YAP (red, right panel) (n = 3). Yellow dotted areas represent nuclear area. Scale bar: 10 μm. scatter plot graph depicting nucleus/cytoplasm ratio of YAP between HCC1806 grown on 0.4 kPa and 20 kPa hydrogels (n=3, n>20). (B) Representative confocal photomicrographs of breast cancer HCC1806 cells untreated (top row of micrographs) or treated with 80 μM TEPP-46, PKM2 cytoplasmic activator (second row from top) grown on 0.4 kPa (soft) hydrogels coated with collagen I and observed after 24 h of culture with staining by DNA (white using DAPI, left panel) and YAP (Red, right panel). Yellow dotted areas represent nuclear area (n = 3). Scale bar: 10 μm. Scatter plot graph depicting nucleus/cytoplasm ratio of YAP between untreated and 80 μM TEPP-46 treated HCC1806 cells on 0.4 kPa hydrogels (n=3, N>25). Error bars denote mean ± SEM. The unpaired Student’s *t* test was performed for statistical significance (**P* ≤ 0.05, ***P* ≤ 0.01, ****P* < 0.001, *****P* < 0.0001).

**Figure S5.** YAP regulates PKM2 localization on stiff matrices in HCC1806. (A) Representative confocal photomicrographs of breast cancer HCC1806 cells untreated (top row of micrographs) or treated with 2μM Verteporfin, YAP signalling inhibitor (second row from top) grown on 20 kPa (stiff) hydrogels coated with collagen I and observed after 24 h of culture with staining by DNA (white using DAPI, left panel) and F-actin (green using phalloidin, right panel (n = 3). Scale bar: 20 μm. Graphs showing cell area (top right) and aspect ratio (bottom) of individual HCC1806 cells untreated and treated with 2 μM Verteporfin on 20 kPa hydrogels (n = 3, N > 30 single cells). (B) Representative confocal photomicrographs of HCC1806 cells untreated (top row of micrographs) or treated with 2 μM Verteporfin (second row from top) grown on 20 kPa (stiff) hydrogels coated with collagen I and observed after 24 h of culture with staining by DNA (white using DAPI, left panel) and PKM2 (Red, right panel) and Scatter plot graph depicting nucleus/cytoplasm ratio of PKM2 between untreated and 2 μM Verteporfin treated HCC1806 cells on 20 kPa hydrogels (n=3, N=50), (n = 3). Scale bar: 20 μm. (C) Epifluorescence photomicrographs of RFP labelled HCC1806 cells with their migration tracks for 3 hours, control (left) and treated with 2 μM Verteporfin (right) on 20 kPa hydrogels and scatter plot graph depicting migration speed (n=3, N > 20). See also Video S15, S16. Error bars denote mean ± SEM. The unpaired Student’s *t* test was performed for statistical significance (**P* ≤ 0.05, ***P* ≤ 0.01, ****P* < 0.001, *****P* < 0.0001).

Video S1: Epifluorescence video of single RFP labelled MDA-MB-231 cells migrating on 0.4 kPa hydrogels, Images were taken from this video for figure 1B

Video S2: Epifluorescence video of single RFP labelled MDA-MB-231 cells migrating on 20 kPa hydrogels, Images were taken from this video for figure 1B

Video S3: Epifluorescence video of single RFP labelled MDA-MB-231 control cells migrating on 0.4 kPa hydrogels, Images were taken from this video for figure 2C

Video S4: Epifluorescence video of single RFP labelled MDA-MB-231 cells treated with PKM2 activator, migrating on 0.4 kPa hydrogels, Images were taken from this video for figure 2C

Video S5: Epifluorescence video of single RFP labelled MDA-MB-231 control cells migrating on 20 kPa hydrogels, Images were taken from this video for figure 3C

Video S6: Epifluorescence video of single RFP labelled MDA-MB-231 cells treated with PKM2 inhibitor, migrating on 20 kPa hydrogels, Images were taken from this video for figure 3C

Video S7: Brightfield video of single MDA-MB-231 control cells migrating on 20 kPa hydrogels, Images were taken from this video for figure 5C

Video S8: Brightfield video of single MDA-MB-231 cells, treated with YAP inhibitor, migrating on 20 kPa hydrogels, Images were taken from this video for figure 5C

Video S9: Epifluorescence video of single GFP labelled HCC1806 cells migrating on 0.4 kPa hydrogels, Images were taken from this video for figure S1C

Video S10: Epifluorescence video of single GFP labelled HCC1806 cells migrating on 20 kPa hydrogels, Images were taken from this video for figure S1C

Video S11: Epifluorescence video of single GFP labelled HCC1806 control cells migrating on 0.4 kPa hydrogels, Images were taken from this video for figure S2C

Video S12: Epifluorescence video of single GFP labelled HCC1806 cells treated with PKM2 activator, migrating on 0.4 kPa hydrogels, Images were taken from this video for figure S2C

Video S13: Epifluorescence video of single GFP labelled HCC1806 control cells migrating on 20 kPa hydrogels, Images were taken from this video for figure S3C

Video S14: Epifluorescence video of single GFP labelled HCC1806 cells treated with PKM2 inhibitor, migrating on 20 kPa hydrogels, Images were taken from this video for figure S3C

Video S15: Brightfield video of single HCC1806 control cells migrating on 20 kPa hydrogels, Images were taken from this video for figure S5C

Video S16: Brightfield video of single HCC1806 cells, treated with YAP inhibitor, migrating on 20 kPa hydrogels, Images were taken from this video for figure S5C

Video S17: Brightfield video of MDA-MB-231 control cells invading into 3D fibrillar matrix. Images were taken from this video for figure 6A, left panel

Video S18: Brightfield video of MDA-MB-231 cells treated with PKM2 inhibitor invading into 3D fibrillar matrix. Images were taken from this video for figure 6A, middle panel

Video S19: Brightfield video of MDA-MB-231 cells treated with YAP inhibitor invading into 3D fibrillar matrix. Images were taken from this video for figure 6A, right panel
