## Supplementary figures and images for "A PKM2-YAP reciprocal repartitioning modulates invasion of breast cancer cells"

### Supplementary figure 1

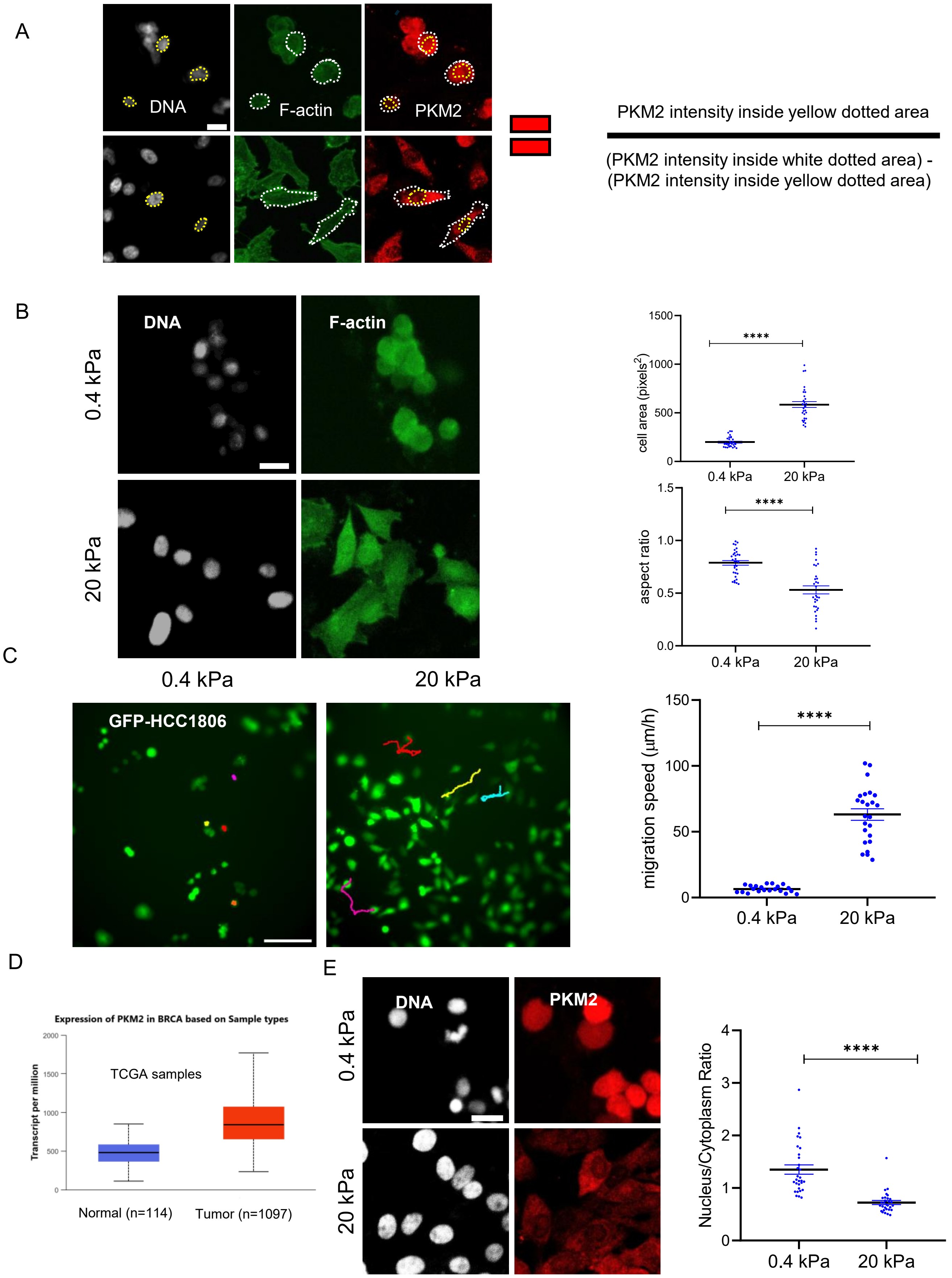

### Supplementary figure 2

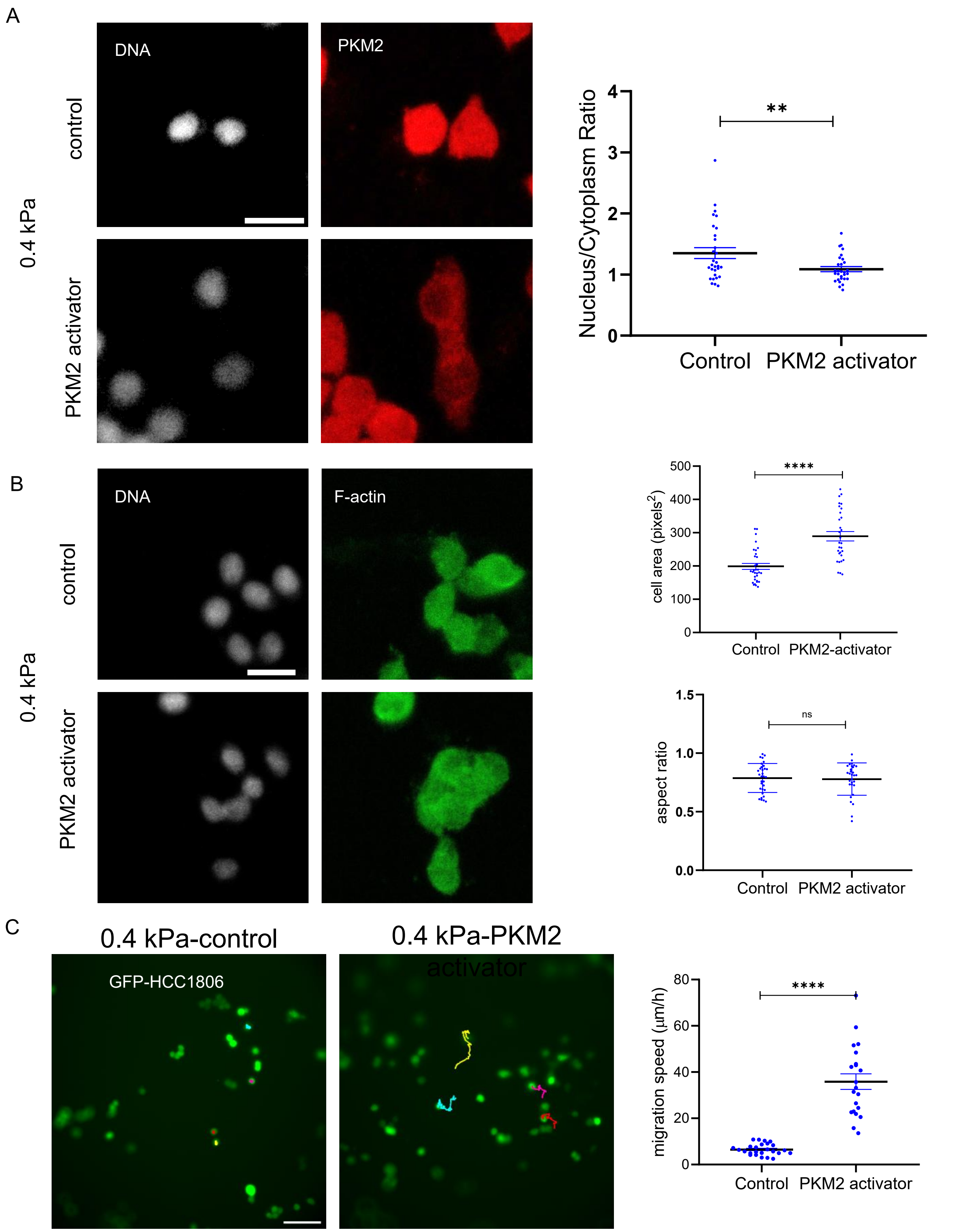

### Supplementary figure 3

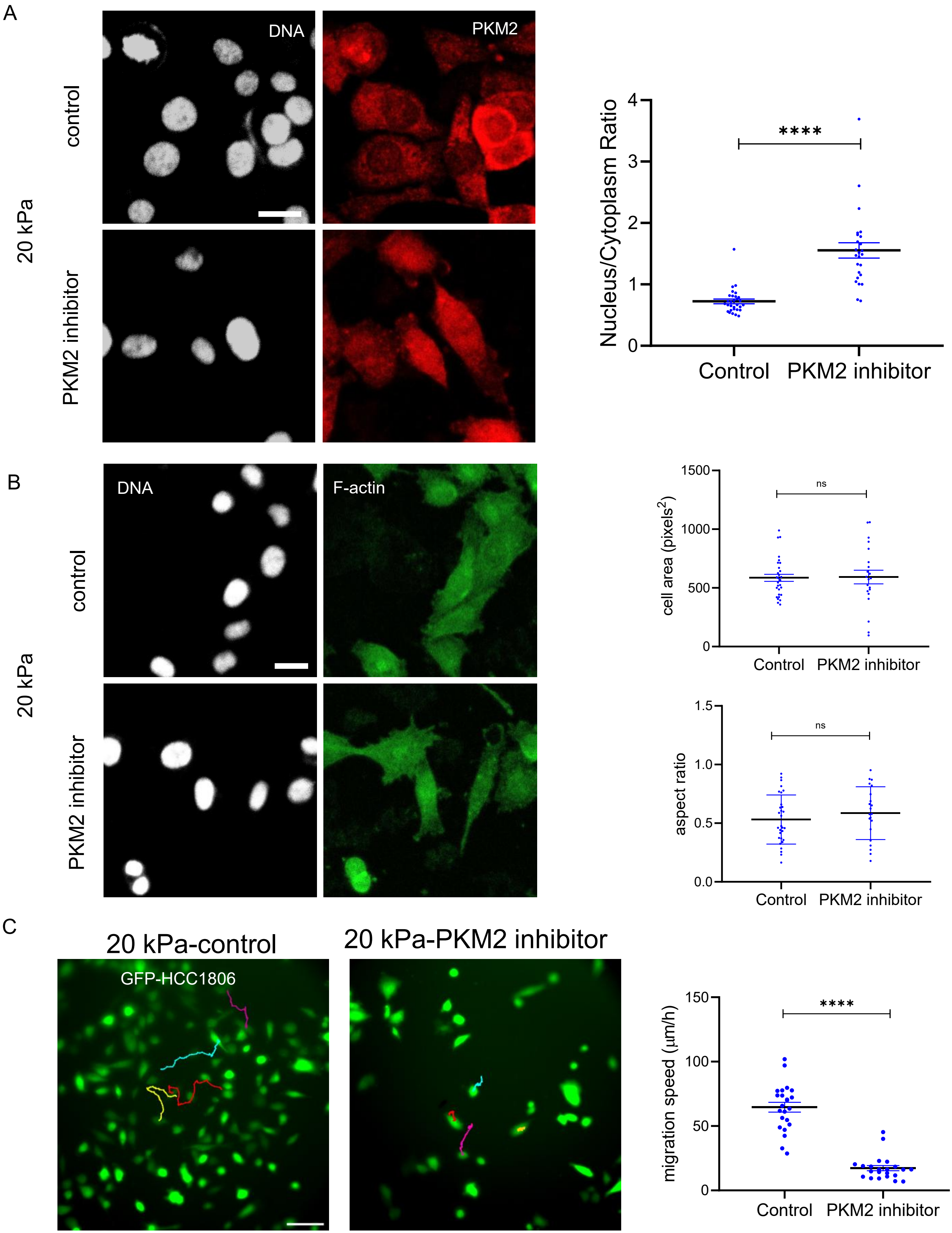

### Supplementary figure 4

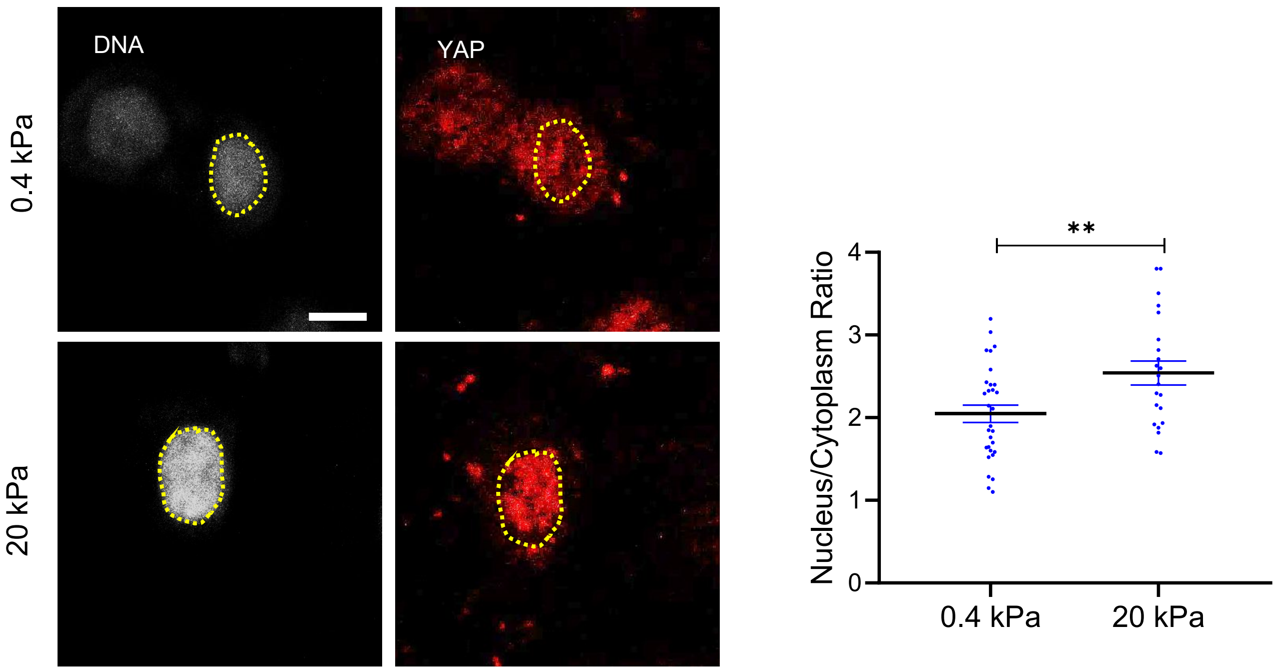

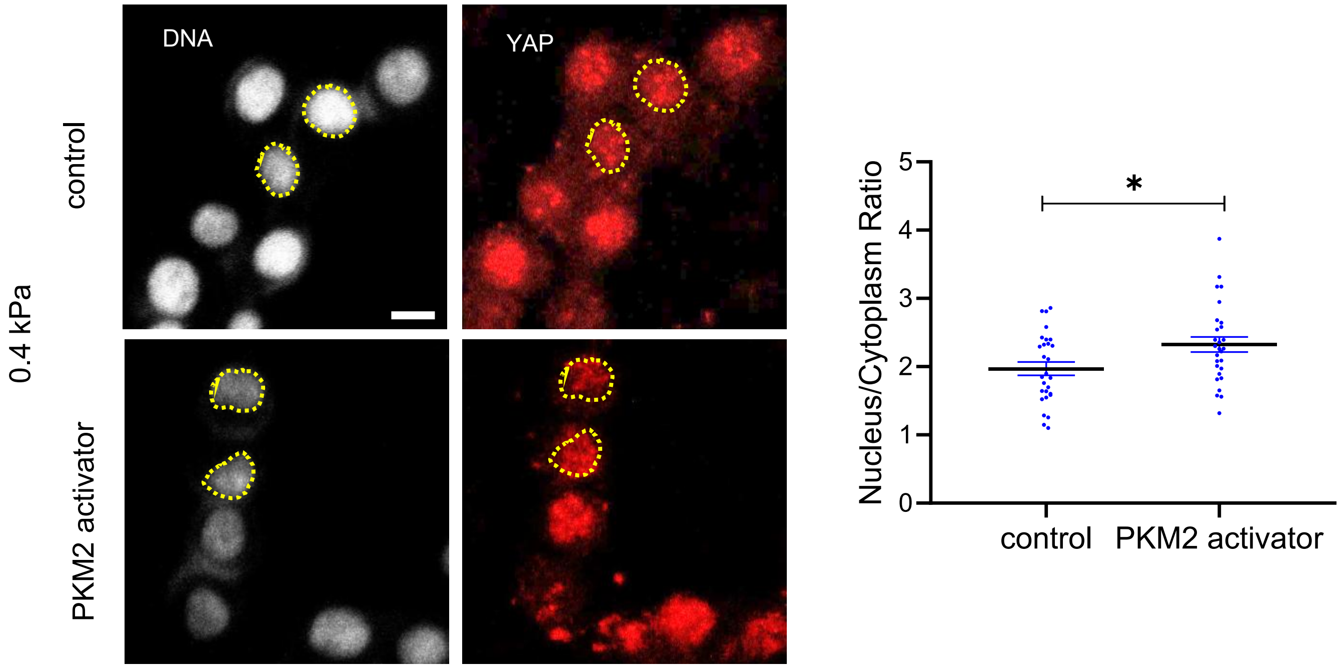

### Supplementary figure 5

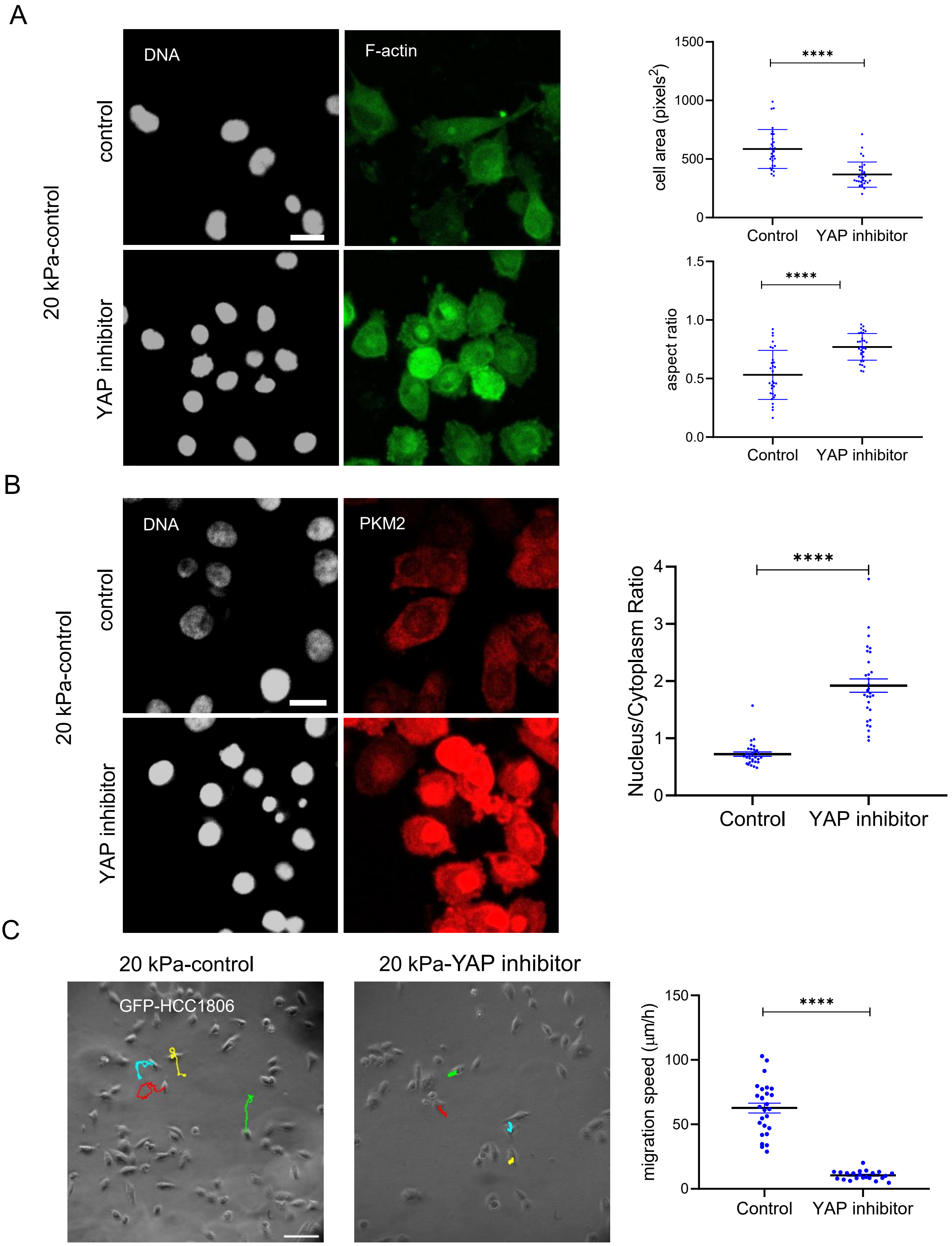
